# Chemical Proteomics Identifies Protein Ligands for Monoacylglycerol Lipids

**DOI:** 10.1101/2024.12.22.629584

**Authors:** Karthik Shanbhag, Amol B. Mhetre, Ojal Saharan, Archit Devarajan, Anisha Rai, M. S. Madhusudhan, Harinath Chakrapani, Siddhesh S. Kamat

## Abstract

Signaling lipids are hormone-like small biomolecules that regulate many critical facets of physiology in mammals, including humans. Given their biomedical importance, the past few decades have seen a tremendous increase in our mechanistic understanding of the physiological processes regulated by a handful of such signaling lipids (e.g.: endocannabinoids, lysophospholipids, prostaglandins). However, a significant number of signaling lipid classes still remain poorly characterized, despite their direct associations to human pathophysiology and disease. Over the past decade, the advent of chemical proteomics technologies coupled with the development of multifunctional lipid probes has rapidly expanded our knowledge in terms of the protein ligands and biological pathways that the different signaling lipids interact with and modulate respectively. While the signaling pathways regulated by the endocannabinoid 2-arachidonoyl-glycerol in mammals are extensively characterized, the same cannot be said for the other members of the monoacylglycerol (MAG) family of signaling lipids. To understand this, here, we report the synthesis of a bifunctional MAG probe, containing a photoreactive group and a biorthogonal handle. Using established chemical proteomics approaches, we profile this bifunctional MAG probe in mouse brain and mammalian cell lysates, and leveraging probe competition experiments identify hitherto unknown protein ligands for MAG lipids. Finally, we biochemically validate the neuronal calcium sensor Hippocalcin as a putative MAG protein ligand, and show for the first time, that MAG may have a role to play in calcium sensing and downstream signaling in the mammalian brain.

## INTRODUCTION

Lipids are indispensable biological building blocks that play vital roles in all forms of life. Amongst the well-known and extensively studied functions of lipids are their ability to maintain structural integrity of cells by forming hydrophobic membranes, and in energy metabolism and storage^1,2^. Besides these two conventional functions, lipids serve as precursors for the biosynthesis of steroidal hormones and have become important biomarkers in the diagnosis of various human diseases, particularly metabolic disorders^1,2^. Lipids have also emerged as important signaling molecules and secondary messengers that modulate diverse biological processes, and over the past two decades, a few signaling lipid classes have been extensively investigated in the context of mammalian physiology^3^. Examples of such well-studied signaling lipids include prostaglandins^4–6^, endocannabinoids^7–10^, and a few classes of lysophospholipids such as sphingosine 1-phosphate^11–15^, lysophosphatidic acid^16–19^ and lysophosphatidylserine^20,21^. Given their importance in mammalian signaling pathways, dysregulation in their metabolism or signaling is linked to numerous human pathophysiological conditions, and drugs targeting their respective metabolic enzymes and cognate receptors are emerging as key therapies in the treatment of an array of human diseases^22,23^.

The endocannabinoid 2-arachidonoyl-glycerol (2-AG, C20:4 MAG) is an endogenous ligand to the cannabinoid receptors in the mammalian brain, and its metabolic and signaling pathways have been very well worked out in the mammalian nervous system^8–10,24^. Given its central role in numerous critical processes in the mammalian nervous system, modulation of 2-AG levels in the brain together with the endocannabinoid system are being rigorously explored as potential therapeutic targets for treating a variety of neurological disorders^25–27^. 2-AG belongs to the monoacylglycerol (MAG) family of signaling lipids^8–10^, and while lot is known with regards to the physiological processes regulated by 2-AG in mammals, the same cannot be said for the other MAG lipids. MAG lipids are biosynthesized from diacylglycerol (DAG) precursors by the action of the DAG lipases, and degraded by MAG lipases (**Figure 1**)^7–10^. Biologically, MAGs are known to exist in two forms, namely 1-MAG (*sn-1* MAG) and 2-MAG (*sn-2* MAG) (**Figure 1**)^28^, and it is speculated that these two forms of MAG can spontaneously (non-enzymatically) interconvert between each other^29^. Besides 2-AG, *in vivo*, MAGs are known to exist as esters of other long chain fatty acids such as palmitic acid (C16:0), oleic acid (C18:1) and linoleic acid (C18:2)^30,31^, and yet very little remains known in terms of the protein ligands and the signaling pathways regulated by these (non 2-AG) MAG variants.

**Figure 1.**
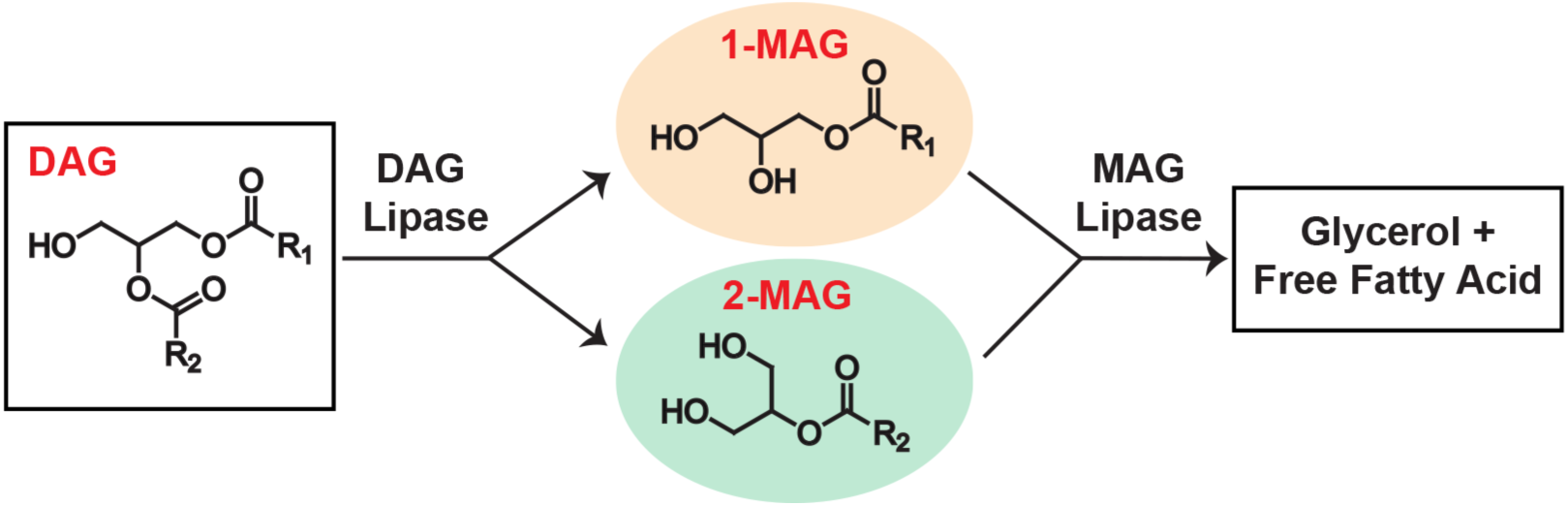
Metabolism of MAG lipids. Monoacylglycerol (1-MAG and 2-MAG) lipids are biosynthesized from diacylglycerol (DAG) precursors by the action of dedicated DAG lipases. On the other hand, they are degraded by the action of MAG lipases to form glycerol and free fatty acids.

Coinciding with our ever-increasing knowledge of signaling lipids, over the past two decades, chemoproteomics (or chemical proteomics) has emerged as a transformative functional proteomics strategy that has enabled the rapid and thorough interrogation of protein-small molecule interactions on a proteome wide scale in complex biological settings^32–34^. Broadly, chemoproteomics approaches can be classified into two different categories, namely activity-based protein profiling (ABPP)^35–40^, and the more recent, photoaffinity based labeling (PAL) strategy^41–45^. Traditionally, ABPP leverages the intrinsic differential reactivity of specific amino acids (such as serine, cysteine, tyrosine) on proteins typically in enzyme active sites or ligand binding pockets^35–40^. On the other hand, the PAL strategy is non-specific in terms of amino acids, and depends almost exclusively on the binding affinity of a particular ligand to protein(s)^41–45^. While ABPP has tremendously aided the discovery and biochemical characterization of enzymes (mostly lipases) involved in the biosynthesis or degradation of signaling lipids^46,47^, the PAL strategy via the development of multifunctional lipid probes has greatly facilitated the identification of novel protein ligands of these signaling lipids, and thus, facilitated our understanding of the biological pathways regulated by them^48^.

In this study, we attempt to use the PAL strategy to identify as-of-yet unknown protein ligands for MAG lipids. Towards this, we develop a bifunctional PAL-compatible MAG probe containing a photoaffinity group and a bioorthogonal handle. Given the abundance of MAG lipids in the mammalian brain, we validate and characterize this MAG probe in the proteomic lysates of the mouse brain and a few mammalian cell lines. Next, leveraging mass spectrometry-based proteomics platforms, and using competition experiments with a control bifunctional free fatty acid probe, we discover novel protein ligands for MAG lipids in the mammalian nervous system. Our modeling and biochemical studies show that the calcium sensor Hippocalcin (exclusively expressed in the central nervous system)^49,50^, is indeed a protein ligand for MAG lipids, and suggests that via an interaction with Hippocalcin, MAG lipids might have a role to play in calcium sensing and signaling pathways in mammalian brain.

## RESULTS

### Development and validation of a MAG probe

In an effort to map the protein ligands of free fatty acids and establish the use of the PAL strategy for doing so, the bifunctional palmitic acid diazirine alkyne (PA-DA) probe (**Figure 2A**) was developed and validated in mammalian cells using various chemoproteomics approaches^51^. We have previously described a facile two-step procedure for the synthesis of 1-MAG lipids^52^, and using the PA-DA probe as the source for the free fatty acid module needed to synthesize 1-MAG lipids, we generated the 1-palmitoyl-glycerol diazirine alkyne (PG-DA) probe (**Figure 2A, Supplementary Figure 1, Supplementary Synthetic Note**) at a milligram scale. Since MAG lipids are quite abundant in the central nervous system, and given our long-standing interest in identifying novel protein interactors for different signaling lipids in the mammalian brain, using various platforms of the PAL strategy (**Figure 2B**)^42,43,48^, we decided to profile the PG-DA probe in proteomic lysates generated from the mouse brain, and a few immortalized mammalian cell lines (Neuro2A, neuronal cell line; BV2, microglial cell lines; RAW264.7, macrophage cell line) that are established surrogates for different cell types in the nervous system.

**Figure 2.**
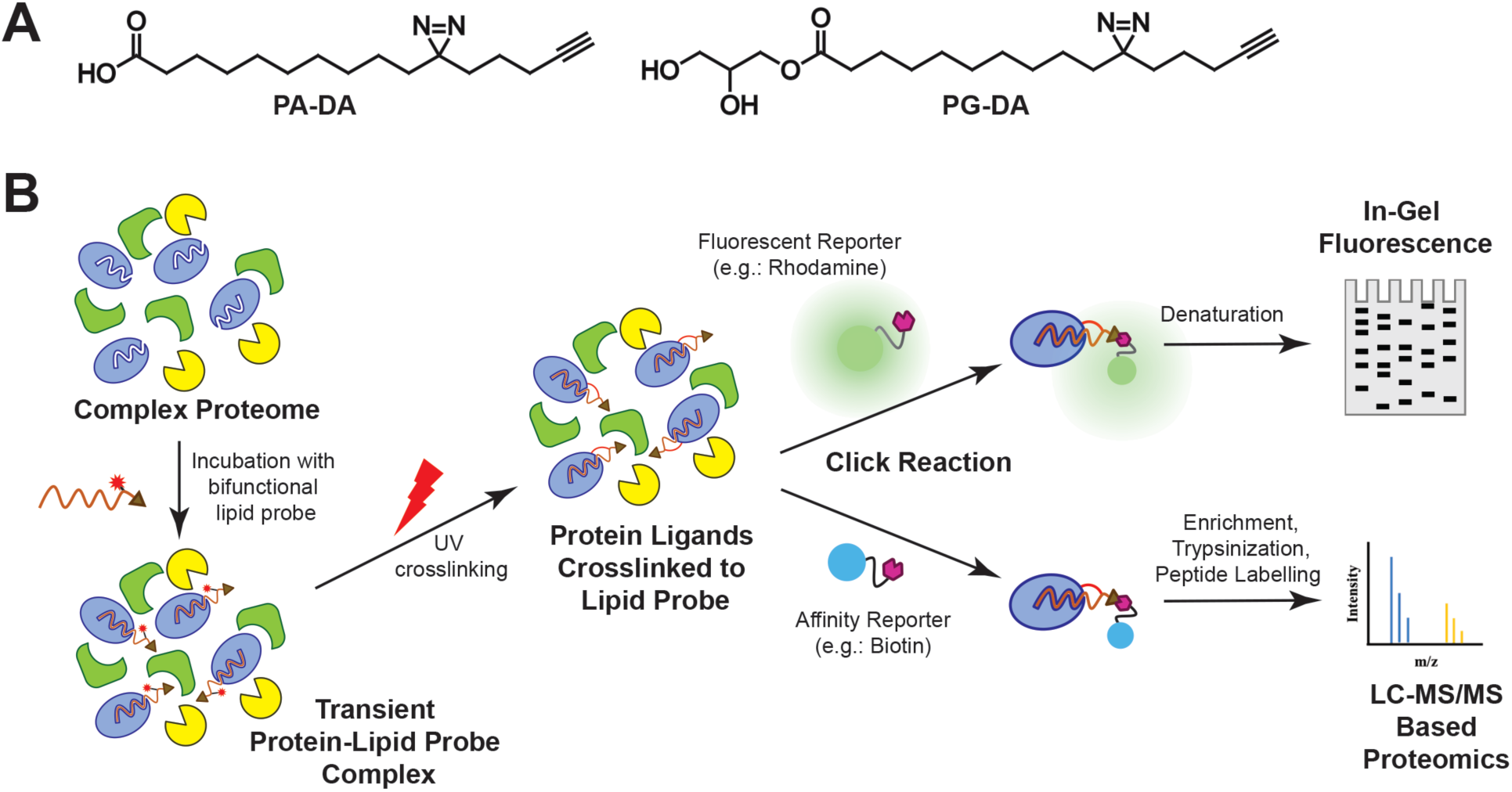
Bifunctional lipid probes and photoaffinity labeling. (**A**) Chemical structures of the palmitic acid-diazarine-alkyne (PA-DA) and 1-palmitoyl-glycerol-diazarine-alkyne (PG-DA) probes used in this study. (**B**) A general workflow for the photoaffinity labeling strategy using both the in-gel fluorescence and the LC-MS/MS based proteomics platforms used in this study.

To validate the photocrosslinking efficacy and downstream enrichment (using click chemistry) of putative protein targets of the PG-DA probe, first, by established gel-based chemoproteomics experiments^53^ with the aforementioned protein lysates, we varied the ultraviolet (UV) crosslinking (irradiation) time (**Supplementary Figure 2**) and probe concentration (**Supplementary Figure 3**) to determine optimal conditions for subsequent experiments with the PG-DA probe. From these gel-based experiments across all the lysates tested, as expected, we found that the photocrosslinking efficacy of the PG-DA probe was UV-dependent, and the optimal probe concentration and UV crosslinking time were found to be 500 μM and 6 mins respectively (**Figure 3A, Supplementary Figure 3**). Since we wanted to use the PA-DA probe as a “control” probe in subsequent competition experiments, we decided to perform similar gel-based chemoproteomics experiments on all lysates using this probe. From this experiment, we found that the photocrosslinking of the PA-DA probe was UV-dependent, and the optimal concentration (**Supplementary Figure 4**) and UV-crosslinking time (**Supplementary Figure 5**) were identical to those of the PG-DA probe.

**Figure 3.**
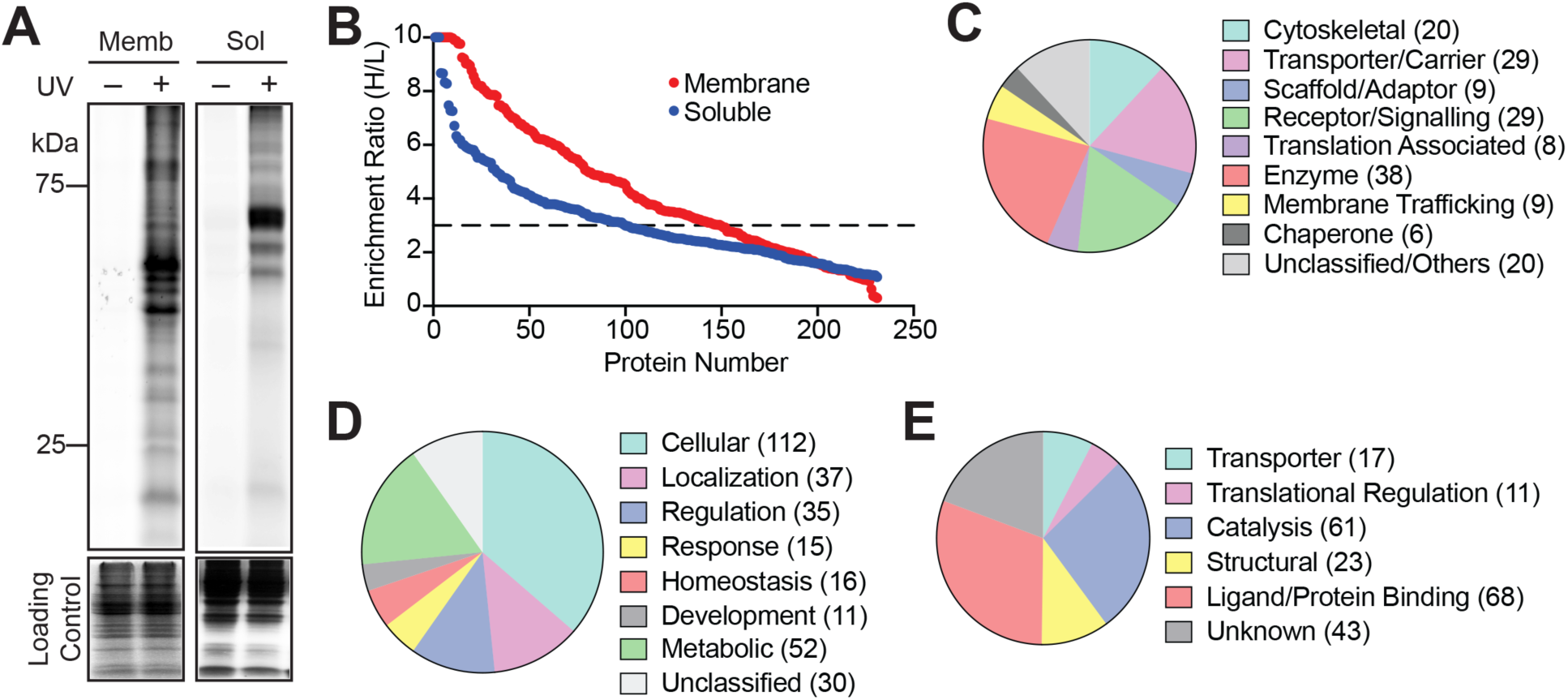
Validation of the PG-DA probe in lysates of the mouse brain. (**A**) A representative fluorescence gel from an in-gel chemical proteomics experiment showing the UV-dependent photocrosslinking of the PG-DA probe (500 μM, 6 mins of UV exposure) to proteins from the membrane (memb) and soluble (sol) lysates prepared from the mouse brain. The Coomassie staining shows the loading control for this experiment. This experiment was done three times with reproducible results each time. (**B**) A LC-MS/MS based chemical proteomics experiment showing enrichment ratio (heavy:light; H:L) of the total proteins identified from the UV-dependent photocrosslinking of the PG-DA probe (500 μM, 6 mins of UV exposure) from the membrane and soluble lysates prepared from the mouse brain. Each data point represents the mean of the enrichment ratio obtained for the respective protein from two or three biological replicate for a particular proteomic fraction, based on the defined filtering criteria for this proteomics experiment. The horizontal dotted line denotes an enrichment ratio ≥ 3, and proteins having an enrichment ratio above this threshold were considered enriched by the PG-DA probe, and taken forward for subsequent analysis. Complete details for all the proteins can be found in **Supplementary Table 1**. (**C–E**) Categorization of proteins enriched by the PG-DA probe from the lysates of the mouse brain based on the Panther database annotation into different: (**C**) protein classes; (**D**) biological processes involved in; and (**E**) known molecular functions.

Having established optimal conditions for assaying the PG-DA probe in different lysates, next, we wanted to identify the repertoire of proteins enriched by it upon UV-crosslinking, leveraging established liquid-chromatography coupled to mass spectrometry (LC-MS) based quantitative chemoproteomics platforms^42,43,48^. In this experiment, the different lysates were treated with the PG-DA probe (500 μM, 6 mins) with or without UV-crosslinking, and the proteins enriched by the PG-DA probe upon photocrosslinking were assessed by LC-MS based quantitative chemoproteomics (**Supplementary Figure 6**). For a protein to be classified as enriched by the PG-DA probe upon UV-crosslinking, it needed to be identified in at least 2 out of 3 replicates, have ≥ 3 quantified peptides per replicate, and have an enrichment ratio ≥ 3 in all replicates. Based on these criteria, in the mouse brain proteomic fractions, we identified a total of 196 proteins (**Figure 3B, Supplementary Table 1**), of which 94 and 46 proteins were unique to the soluble and membrane proteomic fractions respectively, while 56 proteins were identified in both proteomic fractions. Upon further categorization of the proteins enriched by the PG-DA probe from the mouse brain proteomic fraction using the PANTHER database^54,55^, we found that these belonged to different types of proteins (e.g. enzymes, receptors, structural proteins, transporters, adaptors etc.) (**Figure 3C**) and spanned an array of biological pathways and molecular functions (e.g. regulation, signaling, metabolism, development etc.) (**Figure 3D, 3E**). Next, we performed similar quantitative chemoproteomics experiments with the PG-DA probe as a function of UV-crosslinking in the proteomic lysates of different mammalian cells mentioned earlier, and found relative to the results obtained from the mouse brain proteomic fractions, a comparable number of enriched proteins, and similar spread in terms of the types of proteins enriched (**Supplementary Figure 7, Supplementary Table 1**). From the enriched proteins across all these experiments, we find that there are several known lipid interacting proteins, such as transporters, metabolic enzymes, signaling proteins (**Supplementary Table 1**). This result shows that this synthetic bifunctional lipid probe mimics natural lipids, presumably binds to proteins in a ligand-specific manner, and hence, puts the PG-DA probe in good standing towards identifying hitherto unknown protein ligands for MAG lipids in competitive chemoproteomic experiments.

Since we aimed to use the PA-DA probe as a control in subsequent competitive chemoproteomics experiments, we also performed similar quantitative chemoproteomics experiments with this probe as a function of UV-crosslinking in the proteomic fractions of the mouse brain and the mammalian cell lines previously mentioned. Using the same criteria as that of the PG-DA probe, from these experiments, we found that the PA-DA probe was also enrich to a comparable number of proteins without much bias (**Supplementary Figure 8, Supplementary Table 2**), and therefore, this probe could be used at similar concentrations to the PG-DA probe in competitive chemoproteomics experiments towards the identification of MAG protein ligands.

### Competition of PG-DA versus PA-DA probe

To find the protein ligands for MAG lipids, we decided to perform a competitive chemoproteomics experiments using the PG-DA and PA-DA probes (**Supplementary Figure 9**). Here, proteomic lysates from the mouse brain or the aforementioned mammalian cell lines were incubated with the PG-DA (500 μM) or PA-DA (500 μM) probe, following which the UV-crosslinking was performed, and the proteins enriched by the respective probes in different lysates were compared using established quantitative chemoproteomics protocols^42,43,48^. For a protein to be considered for any analysis in this experiment, it needed to be identified in at least 2 out of 3 replicates, and have ≥ 3 quantified peptides per replicate. A protein was considered enriched by the PG-DA probe, if it had an enrichment ratio ≥ 1.5 in all the replicates it was identified, while an enrichment ratio ≤ 0.7 classified a protein to be enriched by the PA-DA probe. Since we were specifically interested in finding novel interactors of MAG lipids, we focused only on the proteins enriched by the PG-DA probe.

Based on the filtering criteria, from the experiments performed on the proteomic lysates from the mouse brain (**Figure 4A**), we found a total of 78 proteins that were enriched by the PG-DA probe, of which, 36 and 33 proteins were found exclusively in the soluble and membrane fraction respectively, while, 9 were found in both fractions (**Supplementary Table 3**). Along similar lines, from the studies performed on the lysates from different mammalian cells, we found that 140, 78, and 179 proteins were enriched by the PG-DA probes from the proteomic fractions of the Neuro2A, BV2 and RAW264.7 cells respectively (**Figure 4B, Supplementary Table 3**). In a quest to identify novel protein interactors of MAG lipids, next, we collated a list of all the proteins that were enriched by the PG-DA probe in the different lysates, and interestingly found, that the PG-DA was able to enrich 364 unique proteins that performed diverse molecular functions (**Figure 4C, Supplementary Table 3**), and were involved in different biological processes (**Figure 4D, Supplementary Table 3**) as per the PANTHER database categorization^54,55^. Specifically, we found that with the exception of the protein Hippocalcin (**Figure 4A**) (expressed exclusively in the mammalian brain, and no other cell line)^49,50^, > 350 proteins were found to be enriched by the PG-DA probe across all the different lysates tested, of which some seemed very consistent across different lysates (**Supplementary Table 3**). Notably, this list of enriched proteins also contained known lipid interactors^53,56,57^ (e.g. Nucleobindin 1 (NUCB1), fatty acid synthetase (FASN), synaptic vesicle membrane protein (VAT1)), thus adding further confidence in the ability of the PG-DA probe to enrich putative protein ligands for MAG lipids.

**Figure 4.**
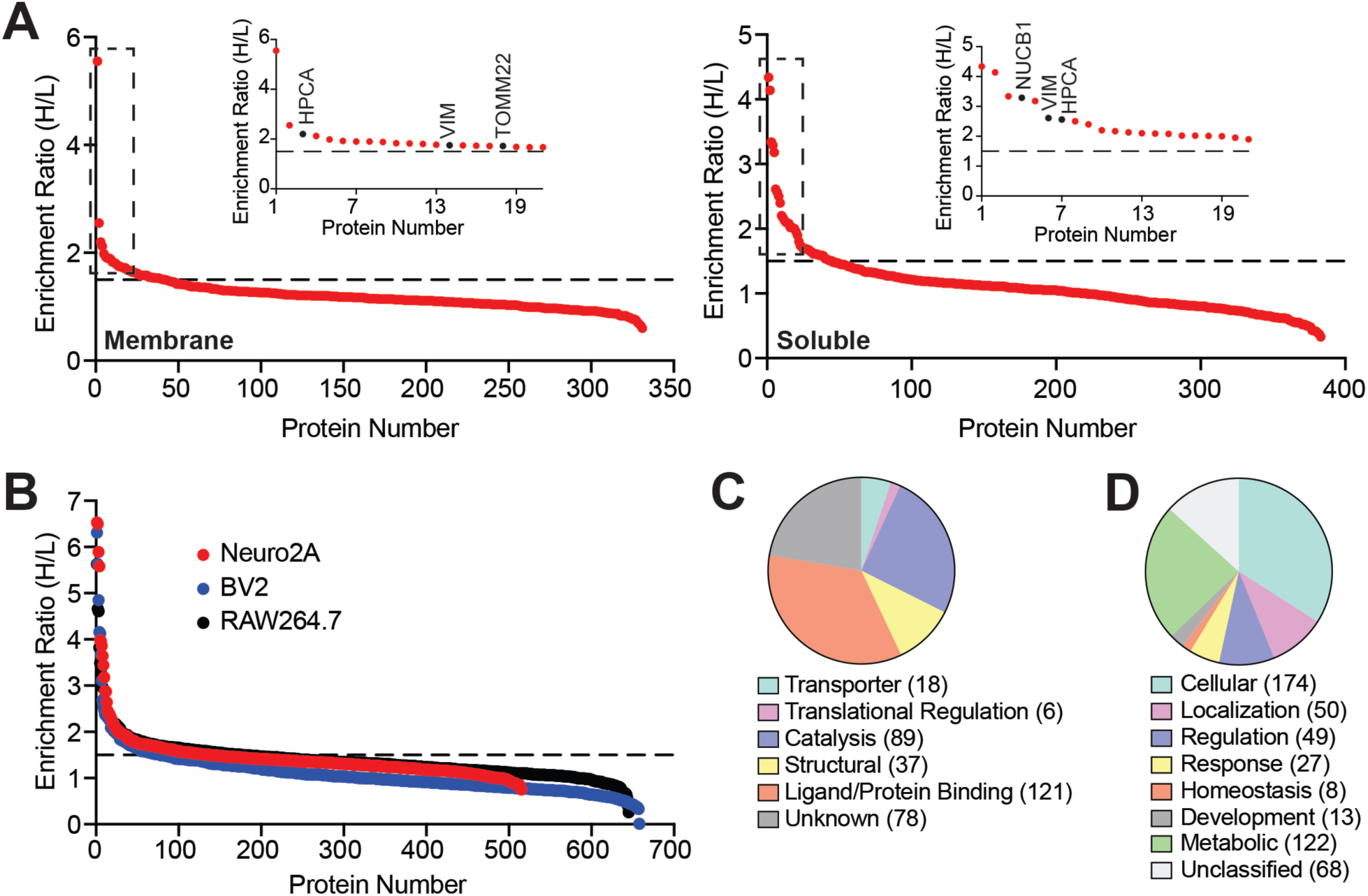
PG-DA versus PA-DA competitive chemical proteomics experiments. (**A, B**) A LC-MS/MS based proteomics analysis showing enrichment ratio (heavy:light; H:L) of the total proteins identified in this probe versus probe (PG-DA versus PA-DA) competitive chemical proteomics experiment (each probe was used at 500 μM, 6 mins of UV exposure) done with (**A**) the membrane and soluble lysates prepared from the mouse brain; and (**B**) lysates prepared from different immortalized mammalian cell lines (Neuro2A, BV2 and RAW264.7). Each data point in (**A**) or (**B**) represents the mean of the enrichment ratio obtained for the respective protein from two or three biological replicate for a particular proteomic fraction, based on the defined filtering criteria for this proteomics experiment. The horizontal dotted line denotes an enrichment ratio ≥ 1.5, and proteins having an enrichment ratio above this threshold were considered specifically enriched by the PG-DA probe, and taken forward for subsequent analysis. Complete details for all the proteins can be found in **Supplementary Table 3**. Inset in (**A**), shows high enrichment of the neuronal calcium sensor Hippocalcin (HPCA), and the mitochondrial translocase sub-unit TOMM22 by the PG-DA probe, along with known MAG interactors such as vimentin (VIM), and nucleobindin 1 (NUCB1). (**C, D**) Categorization of all the proteins specifically enriched by the PG-DA probe based on all experiments described in (**A**) and (**B**), using the Panther database annotation into different: (**C**) known molecular functions; and (**D**) biological processes involved in.

### Molecular docking studies of 1-PG in two putative MAG binding proteins

To validate the outcome of our competitive chemoproteomics experiment, and show that the PG-DA probe was in fact able to enrich protein ligands for MAG lipids, we first decided to use a computational docking strategy to assess if some of the enriched proteins have hydrophobic sites or pockets on them capable of binding the MAG, 1-palmitoyl-glycerol (1-PG). For this experiment, we specifically chose two proteins: the neuronal calcium sensor, Hippocalcin (HPCA)^49,50^, and the translocase of the outer membrane, mitochondrial import receptor subunit 22 (TOMM22)^58–60^. There were two reasons for choosing these proteins: (i) roles of MAG lipids in calcium sensing and mitochondrial import in the brain are poorly understood, and (ii) experimentally elucidated three dimensional structures are available for both these proteins (PDB ID: 5G4P for human HPCA^61^; PDB ID: 7CK6 for human mitochondrial protein translocase that contains TOMM22^62^).

To determine the putative binding site of 1-PG on HPCA and TOMM22, we screened for the presence of hydrophobic pockets that could accommodate a long fatty acyl chain of 1-PG using the CavityPlus^63^ and DEPTH^64^ web servers. Both these algorithms identified nearly identical pockets for each protein (**Supplementary Figure 10**), which were further sorted based on the druggability/ligand binding score, and the cavity with the highest score and largest dimension (≥ 20 Å) was selected for the docking studies (**Supplementary Figure 10, Supplementary Table 4**). Since protein-lipid interactions require hydrophobic surfaces, the electrostatics of these identified pockets were examined (**Supplementary Figure 11**), and both HPCA and TOMM22 showed uncharged (neutral) hydrophobic pockets interspersed with positive or negative charges (presumably to stabilize the head group), capable of 1-PG binding. Next, we attempted to dock 1-PG in the identified hydrophobic pockets of HPCA and TOMM22 using the HADDOCK 2.4 web server^65,66^. Here, the HADDOCK Score which is the weighted sum of all ligand-protein interaction energies, was used to rank and select the best cluster of generated models. For HADDOCK-ing, initially, 10,000 structures were generated for rigid body docking (it0) of which, the top 400 highest-scoring poses underwent semi-flexible refinement (it1). Subsequently, these 400 structures were subjected to final refinement step without a solvent shell (itw). The top 200 structures from this refinement step were then subjected to a RMSD-based clustering using a 1.5 Å cut-off value. Notably, the models within the best clusters for both HPCA and TOMM22, when docked with 1-PG, exhibited remarkably low standard deviations in their HADDOCK scores.

For HPCA, after the final refinement, HADDOCK categorized 183 of the 200 1-PG/HPCA complex models into 3 clusters (**Supplementary Figure 12**). The best scoring cluster amongst these included 25 structures with a HADDOCK score of –35.1 ± 1.5 (RMSD = 0.25 ± 0.15 Å), wherein, 1-PG was docked into the identified hydrophobic pocket. The average van der Waals, electrostatic and ligand binding (ΔG_prediction_) energies of the top four ranked structures in this cluster were –26.4 ± 1.2 kcal/mol, –13.2 ± 2.9 kcal/mol and –8.3 ± 0.1 kcal/mol, respectively (**Supplementary Table 4**). Based on this docking analysis, we predict that 1-PG binds HPCA with the glycerol head situated in a positively charged region near the centre of the identified hydrophobic cavity, and the lipid tail is oriented towards the C-terminal end of the protein (**Figure 5A, Supplementary Figure 12**).

**Figure 5.**
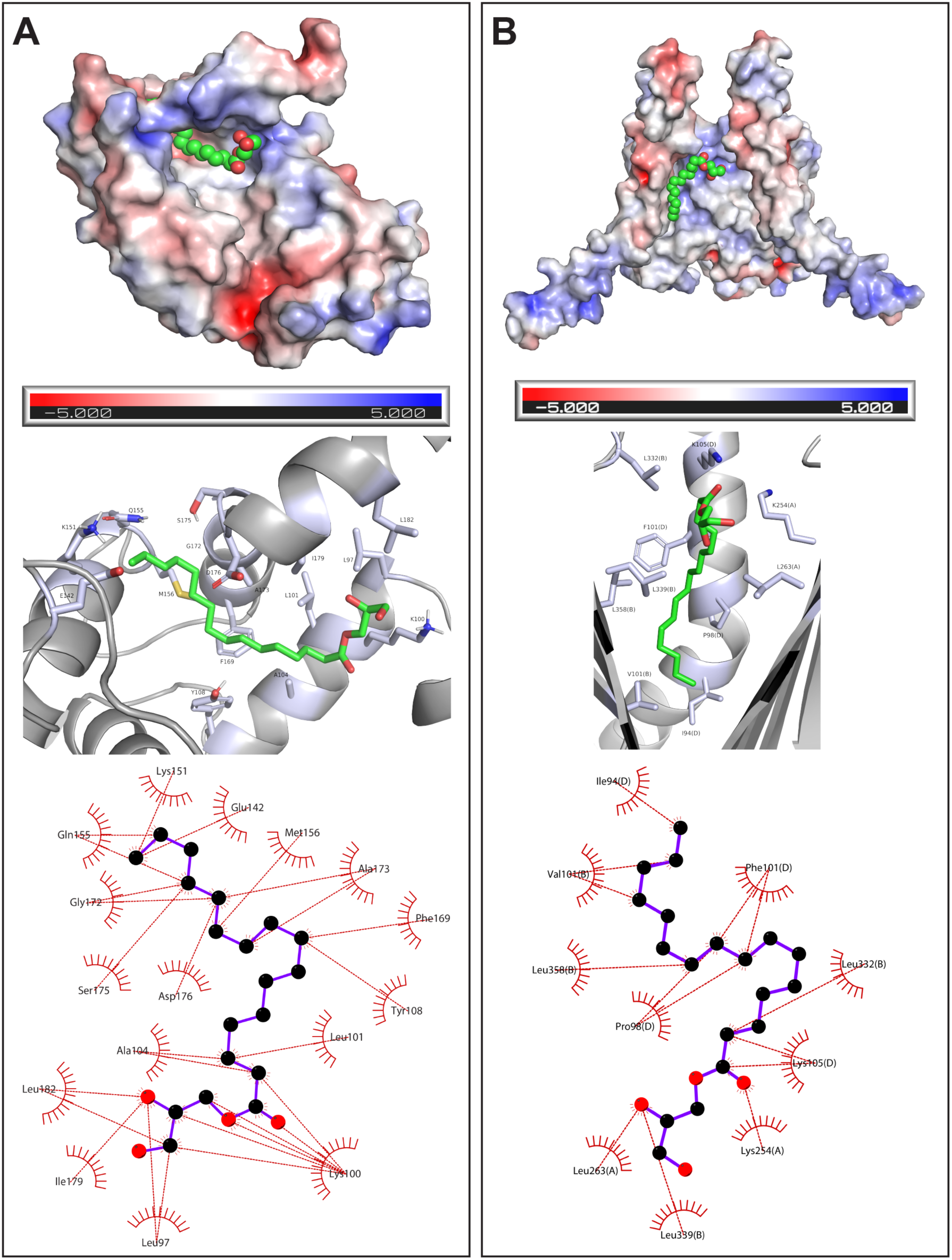
Molecular docking of 1-PG into HPCA and TOMM22. Orientation of 1-PG docked into the identified hydrophobic cavity of (**A**) HPCA and (**B**) TOMM22. In both (**A**) and (**B**), the top panel shows the docking of 1-PG in the hydrophobic cavity of an electrostatic surface model of the respective protein; the middle panel shows the possible residues interacting with 1-PG within this hydrophobic pocket; and the bottom panel shows a two-dimensional rendition illustrating the various residues putatively interacting with 1-PG docked in the hydrophobic pocket.

Similarly, for TOMM22, after the final refinement, 147 of 200 refined models for 1-PG and TOMM22 ligand-protein complex were sorted into 13 clusters. Since HADDOCK does not account for the presence of a lipid bilayer around the mitochondrial translocase complex, the best scoring cluster (HADDOCK Score −37.4 ± 0.5, 8 models clustered) in this case docked 1-PG into the mitochondrial membrane, which is physically impossible. Hence, the second-best cluster, with 1-PG docked into the identified ligand binding pocket with an average HADDOCK score of −35.1 ± 1.1 (RMSD = 0.20 ± 0.13 Å) was selected for further analysis (**Supplementary Figure 13**). This cluster, with 4 of the generated models had average van der Waals, electrostatic and ligand binding (ΔG_prediction_) energies of –19.4 ± 0.8 kcal/mol, –38.1 ± 8.0 kcal/mol and –8.9 ± 0.2 kcal/mol, respectively (**Supplementary Table 4**). 1-PG possibly binds to the same cavity (that phosphatidyl-choline binds to the structure) that opens towards the mitochondrial intermembrane space, with the glycerol head oriented toward the opening of the cavity and the lipid tail entering one of the two possible hydrophobic grooves in the protein complex created by the TOMM40 and TOMM22 proteins (**Figure 5B, Supplementary Figure 13**). The proximity of the lipid tail of 1-PG is towards TOMM22 (to the photoreactive diazirine group of the PG-DA probe also) in the mitochondrial translocase complex, and this possibly explains why TOMM22 is the only component of this multiprotein complex that is enriched by the PG-DA probe.

### Biochemical characterization of HPCA

Having identified hydrophobic pockets in HPCA and TOMM22 capable of binding 1-PG using molecular docking approaches, next, we wanted to validate these findings biochemically. We found from literature that the purification of recombinant TOMM22 was fairly complicated, as this protein was not very stable and needed be purified as part of the entire mitochondrial translocase macromolecular assembly^62^. Hence, we shifted our attention towards biochemically validating HPCA using the PG-DA and PA-DA probes. We successfully expressed mouse HPCA (UniProt ID: P84075) with a N-terminal 6x-His tag in *E. coli*, and were able to highly enrich it (>95%) using affinity chromatography (**Figure 6A**). Next, we incubated the purified mouse HPCA (10 μM) with the PG-DA or PA-DA probe (10 μM, 30 min) at 37 °C, and assessed the ability of these bifunctional lipid probes to bind to purified HPCA as a function of UV-crosslinking (6 min at 37 °C). Consistent with the PG-DA versus PA-DA competition proteomics experiments (**Figure 4A**), we found that the PG-DA bound significantly more to mouse HPCA than the PA-DA probe (**Figure 6B**). Quite surprisingly, we found during this in-gel fluorescence experiment, the lipid probe labeled HPCA (in case of both probes) showed up as two protein bands (at ∼ 20 and 25 kDa) (**Figure 6B**). Upon over-exposing the loading control (Coomassie) gel (**Figure 6B**), we found that HPCA was actually purified in two protein forms: a major form that corresponds to the 20 kDa protein, and a minor form that corresponds to the 25 kDa protein, consistent with previous reports on the purification and biochemical characterization of this protein^61^. Interestingly, we found from this in-gel fluorescence experiment that the minor protein form HPCA (∼ 25 kDa protein band) binds the PG-DA probe far more tightly than the major protein form (∼ 20 kDa protein band) (**Figure 6B**). Next, keeping the concentration of purified mouse HPCA constant (10 μM), we titrated different concentration of the PG-DA or PA-DA probe (0 – 250 μM, 30 min, 37 °C), and found from this dose-response in-gel fluorescence experiment, that at comparable concentrations, the PG-DA probe binds far more to HPCA than the PA-DA probe (**Figure 6C**).

**Figure 6.**
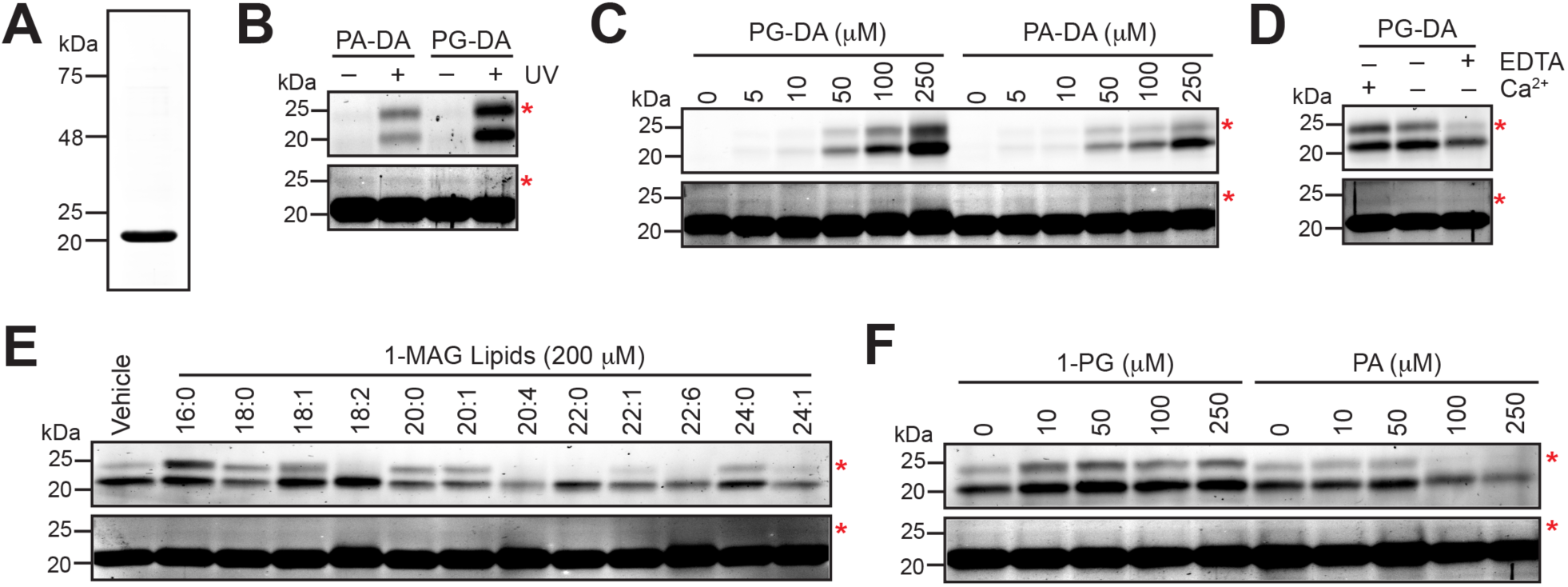
Biochemical characterization of HPCA as a putative MAG protein ligand. (**A**) A representative Coomassie gel showing the enrichment (> 95% purity) of recombinant N-terminal 6x-His tagged mouse HPCA from E. coli using affinity chromatography. (**B**) The UV-dependent preferential binding of the PG-DA probe to recombinantly purified mouse HPCA. In this experiment, the HPCA and lipid probe (PG-DA or PA-DA) concentrations were 10 μM, and 10 μM respectively. (**C**) The dose-dependent binding of the PG-DA probe (0 – 250 μM) or the PA-DA probe (0 – 250 μM) to recombinantly purified mouse HPCA (10 μM). This experiment also shows that at comparable probe concentrations, relative to the PA-DA probe, the PG-DA probe binds better to HPCA. (**D**) The effect of excess calcium (CaCl_2_, 100 μM), or EDTA (100 μM) on the binding of the PG-DA probe (10 μM) to recombinantly purified mouse HPCA (10 μM). (**E**) The effect of incubating different 1-MAG variants (200 μM) on the binding of the PG-DA probe (10 μM) to recombinantly purified mouse HPCA (10 μM). (**F**) The effect of incubating increasing concentrations of 1-PG (0 – 250 μM) or PA (0 – 250 μM) on the binding of the PG-DA probe (10 μM) to recombinantly purified mouse HPCA (10 μM). For all representative gels shown in (**B–F**), the top panel shows the in-gel fluorescence, while the bottom panel is the Coomassie staining as the loading control for that respective gel. For all representative gels shown in (**B–F**), the red asterisk denotes the minor form of HPCA at 25-kDa in both gels. For all representative gels shown in (**B–F**), the Coomassie gel requires over-exposure to visualize this minor band of HPCA. For all gels shown in (**B–F**), the UV exposure time was 6 mins. All experiments shown in this figure (**A–F**), were done at least three times with reproducible results each time.

HPCA is a neuron-specific calcium sensor in the mammalian brain, and we wanted to assess the effect of PG-DA binding to HPCA as a function of calcium concentration. Given its high-affinity for calcium, recombinant HPCA is known to purify in a calcium bound state. Nonetheless, we performed an assay where purified mouse HPCA (10 μM) was first incubated with calcium (100 μM, 30 min, 37 °C) or EDTA (100 μM, 30 min, 37 °C) to chelate any bound calcium. Subsequently. these calcium surplus or devoid forms of HPCA were incubated with the PG-DA probe (10 μM, 30 min, 37 °C), to determine the binding of this lipid probe to HPCA using in-gel fluorescence. From this experiment, we found that relative to the as purified HPCA, the calcium treated HPCA showed marginally more binding to the PG-DA probe than an untreated control (**Figure 6D**). Counter to this, quite interestingly, we found that treating HPCA with EDTA (calcium ion chelator) resulted in a significant loss of PG-DA binding to the protein (**Figure 6D**), suggesting that the binding of 1-MAG lipids to HPCA may be responsive to the calcium sensing function of this neuronal sensor.

Having established the preferential binding of the PG-DA probe to purified HPCA, next, we wanted to assess if incubating HPCA with excess endogenous 1-MAG lipids could compete out this protein-probe interaction, and if there was any structure activity relationship (SAR) for this. For this, we incubated HPCA (10 μM) with a library of 1-MAG lipids (100 μM) previously synthesized by us, together with the PG-DA probe (100 μM, 30 min, 37 °C), and used in-gel fluorescence as a read out for this competitive gel-based chemoproteomics assay. From this experiment, we found that the very-long chain fatty acid containing 1-MAG variants (chain length ≥ C20), especially those having polyunsaturated fatty acids (e.g. C20:4, C22:6) were able to efficiently compete the binding of the PG-DA probe to HPCA (**Figure 6E**). On the contrary, we found that the incubation of HPCA with a few long-chain fatty acid containing 1-MAG lipids (C16:0, C18:1), resulted in the increased binding of the PG-DA probe to HPCA (**Figure 6E**). To test this premise, we incubated purified HPCA with increasing concentrations of 1-PG or palmitic acid (PA) (0 – 250 μM) in the presence of the PG-DA probe (10 μM, 30 min, 37 °C) to assess effect of 1-PG or PA on binding of this lipid probe to HPCA using in-gel fluorescence as a readout. Consistent with the previous SAR experiment, we found that increasing the 1-PG concentration resulted in the heightened binding of the PG-DA probe (in a dose-dependent manner from 0 – 50 μM, and constant from 50 – 250 μM) (**Figure 6F**). Counter to this, we found that incubation of HPCA with PA resulted in a dose-dependent reduction in the binding of the PG-DA probe to the protein (**Figure 6F**). Taken together, our results from the biochemical characterization of this neuronal sensor collectively suggest that there must be an in-built selectivity for HPCA in recognizing and binding 1-MAG lipids, which seems to be highly cooperative in nature, and somehow linked to the calcium sensing/binding function of this protein.

## DISCUSSION

Over the past two decades, chemical proteomics has emerged as an invaluable tool in mapping the physiological protein ligands to an array of small bioactive molecules such as drugs, xenobiotics, and endogenous signaling molecules like hormones, and signaling lipids^32–34^. In particular, the development of synthetic strategies towards making multi-functional lipid probes in tandem with the versatility of bioorthogonal reactions has seen a steep rise in our annotation of protein ligands to a number of signaling lipids, and unraveled interesting biological pathways regulated by them^48^. Amongst the signaling lipids, given their biomedical importance, the endocannabinoids have been extensively investigated over the years. 2-AG, an endocannabinoid, belonging to the MAG family remains the most studied member of this class of signaling lipids, mostly in the context of the nervous system^8–10^. Yet interestingly, besides 2-AG, the role that other MAG lipids play in mammalian physiology and the proteins that they interact with remain cryptic. Recent studies have however shown that abundant non-2-AG MAG variants (e.g. C16:0 MAG, C18:1 MAG) have an important role to play in glucose stimulated insulin signaling^67,68^, and in the regulation of food-satiety index^69^. However, the precise mechanisms, their protein-partners, and the signaling pathways modulated by these abundant non-2-AG MAG variants remain unknown in this physiological context.

To map the putative protein ligands for non-2-AG MAG lipids, here, we report the synthesis of the PG-DA probe. Initially, using in-gel fluorescence as a readout, and leveraging established chemical proteomics techniques, we successfully validate the UV-dependent photocrosslinking efficiency of this bifunctional C16:0 MAG probe, along with the previously reported PA-DA probe, in lysates from the mouse brain and different immortalized mammalian cell lines (Neuro2A, RAW264.7 and BV2) in an effort to standardize downstream protocols to use PG-DA in tandem with the PA-DA probe to identify as-of-yet unknown protein interactors of MAG lipids. Next, using established LC-MS/MS based chemical proteomics platforms with the aforementioned lysates, we show that the PG-DA can enrich a significant number of proteins that belong to diverse protein families, have a variety of physiological functions and are involved in different biological pathways. Further, from exhaustive competition experiments of the PG-DA probe versus PA-DA probe using advanced LC-MS/MS based chemical proteomics platforms, we identify and report new protein interactors of MAG lipids in the mammalian nervous system. From these novel MAG protein ligands, we handpicked TOMM22 and HPCA, as putative MAG-binding proteins to validate using a molecular docking approach. Our *in-silico* analysis showed that both TOMM22 and HPCA possess deep hydrophobic pockets (or cavities), where 1-PG can bind in an energetically favorable conformation. Lastly, we recombinantly purify HPCA, and show that this neuronal calcium sensor has a strong preference to bind 1-MAG lipids. Our studies also suggest that HPCA’s interaction with 1-MAG lipids likely influences its ability to bind calcium or vice versa, and that this protein perhaps has an in-built selectivity and cooperativity towards binding different 1-MAG variants.

Under basal physiological conditions HPCA is exclusively expressed in the mammalian brain, where it functions as a calcium sensor in neurons, and is predominantly localized to the cytosol^49,50^. During synaptic activation/events, the excitation of the NMDA receptor causes an influx of calcium into neurons^70–72^. This increased intracellular calcium concentration is sensed by neuronal HPCA via its EF-hand structural domains^73^, and results in the myristoylation of HPCA^74,75^. This myristoylated form of HPCA then translocates to the plasma membrane, interacts with the membrane anchored AP2-adaptor complex, and promotes the endocytosis of the counter AMPA receptor to potentiate memory formation via signaling from the activation of the NMDA receptor^49,50^. Interestingly, another study along similar lines shows that the enzymatic activity ABHD6 (a putative MAG lipase) is correlated to MAG levels in the brain, and this ABHD6 activity in turn, influences the endocytosis of AMPA receptors^76^. This literature precedence, together with our biochemical studies, strongly suggest that there must exist a functional crosstalk between the calcium sensing activity and MAG-binding ability of HPCA, and these cooperatively are modulating the signaling events between the NMDA-AMPA receptors in the mammalian brain. Moving forward, it will be useful to understand these neuronal mechanisms in the context of HPCA, and perhaps leverage the use of 1-MAG variants (or similar compounds) as pharmacological tools in dissecting out the detailed steps involved in this process.

Besides HPCA, we also show using *in-silico* analysis that TOMM22 binds 1-PG. Given this, moving forward it will be interesting to see if and how 1-MAG variants are involved in different mitochondrial functions such as respiration. Additionally, our competitive proteomics experiments also identify quite a few hitherto unknown 1-MAG protein interactors, and for several of these identified proteins, a biochemical function or endogenous protein ligand is lacking. Thus, our data opens up several new research avenues involving MAG lipids, and the tools and probes described here, can facilitate the discovery of novel signaling pathways modulated by MAG lipids in tissues other than the mammalian brain. Finally, the lipid probes described here, might also prove useful tool compounds in expanding the identification of hydrophobic pockets on protein or protein-protein interfaces, and these could be exploited as potential druggable hotspots for developing new therapeutics for an array of human diseases.

## MATERIALS AND METHODS

### Materials

Unless mentioned otherwise, all chemicals and reagents were purchased from Sigma-Aldrich, and all tissue culture media and consumables were purchased from HiMedia.

### Mice brain harvesting

All experiments involving mice used in this study were approved by the Institutional Animal Ethics Committee at IISER Pune constituted as per the guidelines provided by the Committee for the Purpose of Control and Supervision of Experiments on Animals, Government of India (Protocol No.: IISER_Pune IAEC/2023_03/02). All mice used in this study were males from the C57Bl/6J strain, 10 – 12 weeks of age, and housed at the National Facility for Gene Function in Health and Disease (NFGFHD) at IISER Pune. In this study, the mice were deeply anaesthetized using isoflurane, and euthanized by cervical dislocation. Subsequently, their brains were surgically harvested, split sagittally into two equal anatomical parts, washed 2x with cold sterile Dulbecco’s Phosphate Buffered Saline (DPBS), transferred into a 1.5 mL microcentrifuge tube, flash frozen using liquid nitrogen, and stored at –80 °C until further use.

### Mammalian cell culture

All immortalized mammalian cell lines used in this study, namely Neuro2A, RAW264.7 and BV2, were purchased from ATCC, and cultured in high glucose Dulbecco’s Modified Eagle Medium (DMEM) (HiMedia; Catalog No: AL066A) supplemented with 10% (v/v) Fetal Bovine Serum (FBS) (HiMedia; Catalog No: RM1112) and 1x Penicillin-Streptomycin (HiMedia; Catalog No: A001A) at 37 °C with 5% (v/v) CO_2_. To prevent activation of RAW264.7 and BV2 cells, the FBS used in this media was heat inactivated by heating at 60 °C for 45 mins. All cells were routinely stained with 4’,6-diamidino-2-phenylindole (DAPI) using previously described protocols^77,78^, to ensure that they were devoid of any mycoplasma contamination. All cells were grown to at least 70% confluence, after which they were harvested by scraping, washed 3x with cold sterile DPBS, pelleted in a 1.5 mL microcentrifuge tube, flash frozen in liquid nitrogen, and stored at –80 °C until further use.

### Preparation of proteomic lysates

Proteomic lysates from the mouse brain and mammalian cells were prepared using previously described protocols^52,77,78^ in sterile DPBS, and their concentrations were estimated using the Bradford assay^79^. To specially ensure that the PG-DA probe was intact in our experiments, the proteomic lysates were pre-treated with 2 mM phenylmethylsulfonyl fluoride (PMSF) for 45 mins at 37 °C to inactivate resident lipases that might potentially hydrolyze the PG-DA probe.

### In-gel fluorescence studies

For a typical in-gel fluorescence experiment, 100 μL of the lysate (concentration 2 mg/mL) was incubated with the PG-DA or PA-DA probe (probe concentration and time of incubation mentioned in the gels represented in the figures as per the experiment) at 37 °C with constant shaking in the dark, following which the UV-crosslinking step at 365 nm (UV-crosslinking time mentioned in the gels represented in the figures as per the experiment) was performed. After UV-crosslinking of the PG-DA or PA-DA probe, and the lysates were incubated with 11 μL of the Click reaction mixture [6 μL of Tris-(benzyltriazolylmethyl)-amine (TBTA) (Sigma; Catalog No: 678937-50MG) (1.7 mM in 4:1 DMSO: *tert*-butanol) + 2 μL CuSO_4_ (50 mM in MilliQ water) (Avra Synthesis Pvt. Ltd.; Catalog No: ASC1746) + 2 μL Tris-(2-carboxyethyl)-phosphine (TCEP) (50 mM, freshly prepared in MilliQ water) (Sigma; Catalog No: C4706-2G) + 1 μL rhodamine-azide (10 mM in DMSO) (Sigma; Catalog No: 760765-5MG)] for 60 mins at 25 °C with constant shaking. The Click reaction was quenched by adding 33 μL of 6x SDS-PAGE loading buffer, and the samples were resolved on a 10% SDS-PAGE gel. The gel was visualized for in-gel fluorescence from probe crosslinking on an iBright 1500 gel imager (ThermoFisher Scientific).

### Proteomics sample preparation and LC-MS analysis

All proteomics samples were prepared using the lysates that were processed as described earlier. For a typical proteomics experiment, 1 mL of the lysate (concentration 2 mg/mL) was incubated with the PG-DA probe (500 μM) or the PA-DA probe (500 μM) for 30 mins at 37 °C with constant shaking in the dark. Following this, the UV-crosslinking step (UV-crosslinking time mentioned as per the experiment later) was performed, and the probe treated lysates were incubated with 110 μL of the Click reaction mixture [60 μL of TBTA (1.7 mM in 4:1 DMSO: *tert*-butanol) + 20 μL CuSO_4_ (50 mM in MilliQ water) + 20 μL TCEP (50 mM, freshly prepared in MilliQ water) + 10 μL biotin-azide (10 mM in DMSO) (Sigma; Catalog No: 762024-25MG)] for 60 mins at 25 °C with constant shaking. Thereafter, the lysates were denatured, reductively alkylated with iodoacetamide and digested with proteomics grade trypsin (Promega; Catalog No.: V5111) using protocols previously reported by us^77,80^. For the quantitative proteomics experiments, we used the established reductive demethylation (ReDiMe) peptide labeling strategy previously described by us^77^. In the experiments involving lipid probes in the ± UV crosslinking study, the tryptic peptides obtained from the UV-crosslinking group were labeled with heavy formaldehyde (CD_2_O) (Cambridge Isotope Laboratories Inc.; Catalog No: DLM-805-20), while those from the no UV group were labeled with light formaldehyde (CH_2_O) (Sigma; Catalog No: 252549-25ML). On similar lines, in the experiments comparing the targets from PG-DA versus PA-DA, the tryptic peptides obtained from the PG-DA group were labeled with heavy formaldehyde, while those from the PA-DA group were labeled with light formaldehyde. After the ReDiMe labeling for the respective experiment, the heavy and light labeled peptides were mixed, and desalted using the established StageTip protocol^81^. All LC-MS/MS was performed on a Sciex TripleTOF6600 mass spectrometer fitted with a front-end Eksigent nano-LC 425 system using LC columns and run conditions previously reported by us^80^. Briefly, all proteomic samples were acquired using an information-dependent acquisition (IDA) mode over a *m/z* = 200 – 2000. In our experiments, a full MS survey scan was followed by the MS/MS fragmentation of the 15 most intense peptides. In all our experiments, a dynamic exclusion was also enabled (repeat count, 2; exclusion duration, 6 sec) to increase peptide coverage. Peptide identification and quantification was carried out using Protein Pilot (version 2.0.1, Sciex) using the in-built Pro-Group and Paragon algorithms against the RefSeq protein database of *Mus musculus* (Release 109, last modified on 22^nd^ September 2020) generated by UniProt. While searching the peptides, we defined iodoacetamide alkylation of cysteine as a static modification and the oxidation of methionine and N-terminal acetylation as variable modifications. The ReDiMe algorithm was selected within Protein Pilot software for quantification of identified proteins. In all our proteomic searches, the precursor ion and MS/MS mass tolerance were set at 20 and 50 ppm, respectively, for the peptide searches. Additionally, the false discovery rate (FDR) was calculated using a decoy database search, and a stringent FDR < 1% was used to filter proteins and peptides in our experiments. In all the analysis reported in this study, a maximal cut-off ratio of 10 was imposed on the enrichment ratio (heavy label to light label) for the ReDiMe labelling experiments.

### Molecular docking of 1-PG into HPCA and TOMM22

The three-dimensional structures of HPCA (PDB ID: 5G4P, UniProt: P84074) and the TOMM22 complex (PDB ID: 7CK6, UniProt: Q9NS69) were obtained from the Protein Data Bank^82^. The calcium ions from HPCA and the 1,2-distearoylphosphatidylcholine from the TOMM22 complex were removed prior to docking. Further, the TOMM22 complex was also amended to appear as a single chain for docking using the PDB-Tools web server^83^. Additionally, the coordinates for 1-PG [IUPAC: (2S)-2,3-dihydroxypropyl hexadecanoate] were also obtained from the PDB. The potential ligand binding sites in HPCA and the TOMM22 complex were identified using the CavityPlus^63^ and DEPTH^64^ web servers with their default settings, and 1-PG was docked onto them using the HADDOCK2.4 web server^65,66^. Briefly, residues identified from cavity predictions were supplied to HADDOCK to define ambiguous interactions restraints (AIRs). These AIRs, input as active and passive residues, were defined as amino acid residues whose side chains point toward the identified cavity. The list of residues used as restraints for the two proteins/complexes are as follows: for HPCA: *Active residues - K151, Q155, S175; Passive residues - Y52, F64, T92, S93, W103, A104, Y108, I128, R148, G172, A173, D176*; and for the TOMM22 complex: *Active residues - K330/TOM40(A), K105(C), K105(D); Passive residues - W86(C), P98(C), E102(C), K105(C)*. The ligand (1-PG) was defined as a fully flexible active residue. 10,000 structures were generated for rigid body docking (it0) of which 400 highest-scoring poses were selected for semi-flexible refinement (it1). A 7Å cut-off was used for defining flexible regions at the interface of ligand-protein contact. These 400 structures were then subject to final refinement without a solvent shell, and the top 200 structures were used for RMSD-based clustering with a cut-off value of 1.5Å. Default values were used for all other parameters for protein-ligand docking. Clusters were manually inspected for ligand binding in the identified pocket, and all structures were visualized using the PyMOL software. The surface electrostatics and electrostatic potentials for HPCA and the TOMM22 complex were determined using the APBS plugin for PyMOL using the AMBER force field^84^. The ligand binding energies (ΔG_prediction_) of 1-PG to HPCA and the TOMM22 complex were calculated using the PRODIGY-LIG web server^85^, and 2D pharmacophores were visualized using LigPlot+^86^.

### Purification of recombinant HPCA

The mouse HPCA cDNA (Horizon Discovery; Catalog No: MMM1013-202769681) was cloned into the pET45b(+) vector, between the BamHI and XhoI restriction sites, such that the protein is eventually expressed as with a N-terminal 6x-His tag. The resulting plasmid was sequenced to ensure that the gene of interest (HPCA) was in the correct reading frame, and was transformed into E. coli BL21 (DE3) competent cells. A single colony was picked and grown overnight in 10 mL Luria-Bertani (LB) media containing ampicillin (final concentration of 100 μg/mL) at 37 °C with constant shaking. This primary culture was used to inoculate 1 L of the same medium, and the cells were subsequently grown at 37 °C with constant shaking (∼ 180 rpm) till the OD_600_ reached ∼ 0.6. At this point, the protein expression was induced by adding 1 mM isopropyl β-D-1-thiogalactopyranoside (IPTG) and the cells were grown overnight at 20 °C with constant shaking. Overexpression was confirmed SDS-PAGE analysis, and the cells were harvested by centrifugation at 6000g for 20 mins at 4 °C. The resulting cell pellet (∼ 3 g/L culture) was re-suspended in 50 mL of lysis buffer (50 mM Tris.HCl, 150 mM NaCl, 10 mM imidazole, 2 mM PMSF at pH 8.0), and lysed using a probe sonicator for 20 min with 10 sec ON/OFF cycle with 60% amplitude. The resulting lysate was centrifuged at 30,000g for 30 mins at 4 °C, and the supernatant was applied to a HisTrap FF column (Cytiva; Catalog No: 17525501) which was pre-equilibrated with the binding buffer (50 mM Tris.HCl, 150 mM NaCl, 10 mM imidazole at pH 8.0) to enrich 6x-His tagged HPCA. The protein of interest (6x-His tag HPCA) was eluted from the column using an increasing gradient of imidazole (50 to 250 mM) as per the manufacturer’s instructions. The collected protein fractions were assessed by SDS-PAGE analysis, and those containing the 6x-His tagged HPCA were pooled together, and dialyzed overnight at 4 °C in 50 mM Tris.HCl (pH 8.0) using a 10-kDa molecular weight cut-off membrane (ThermoFisher; Catalog No: 68100) to get rid of excess imidazole. The resulting protein was concentrated to a final concentration of 25 mg/mL (using an Amicon Ultra-15 Centrifugal Unit, Millipore; Catalog No.: UFC901024), flash frozen using liquid nitrogen as 10 μL aliquots, and stored at –80 °C until further use. Typical protein yields were 12.5 mg of purified 6x-His tagged HPCA/L culture. All in-gel fluorescence studies 10 μM of protein was used in a final volume of 100 μL in the assay buffer (50 mM Tris.HCl at pH 8.0). All UV-crosslinking step, Click reaction and its quenching were the same as described earlier. All in-gel fluorescence studies using HPCA was performed on a 12.5 % SDS-PAGE gel, and imaged as described earlier.

### Synthesis of PA-DA and PG-DA

Complete details of the synthesis of PA-DA and PG-DA, along with compound characterization data can be found in the **Supplementary Synthetic Note**.

## Supporting information

Supplementary Synthetic Note

## ACKNOWLEDGEMENTS

We acknowledge financial support from the Science and Engineering Research Board (SERB), Department of Science and Technology, Government of India (Grants: SB/SJF/2021-22/01 to S.S.K.; and CRG/2023/003892 to H.C.), an EMBO Young Investigator Award (to S.S.K.), Department of Biotechnology, Government of India under the BICB Centre Scheme (Grant: BT/PR40262/BTIS/137/38/2022 to IISER Pune), and the Prime Minister’s Research Fellowship (graduate student fellowship to K.S. and O.S.). We thank Saddam Shekh for technical assistance and maintenance of the biological mass spectrometry facility at IISER Pune. We thank Pooja Thakral for providing the PA-DA probe that was used in the studies reported in Figure 6B and 6C. All staff members of the NFGFHD at IISER Pune (supported by a grant from the Department of Biotechnology, Govt. of India; BT/INF/22/SP17358/2016) are thanked for maintaining and providing mice for this study. Members of the S.S.K. and H.C. labs at IISER Pune are thanked for providing critical comments and inputs throughout the course of this study.

## AUTHOR CONTRIBUTION

K.S. performed all the chemical proteomics studies with assistance from O.S.; K.S. performed all the biochemical studies with assistance from A.R.; A.D. performed all the molecular docking studies with inputs from M.S.M., K.S., and S.S.K.; A.M., H.C. and S.S.K. designed the synthetic route for making the chemical probes; A.M. developed the synthetic methodology for the chemical probes; H.C., and S.S.K. conceived the project, supervised this project and acquired funding for this project. K.S., H.C., and S.S.K. wrote the manuscript with inputs from all authors.

## DATA AVAILABLITY

All data that supports the findings of this study are available in the paper and its associated Supporting Information or are available with the Lead Contact upon reasonable request.

All raw data from the proteomics experiments are available on the PRIDE database with accession numbers: PXD051332 (PG-DA probe ± UV), PXD051775 (PA-DA probe ± UV), and PXD051399 (PG-DA probe vs PA-DA probe competition).

## CONLFICT OF INTEREST

The authors declare no conflict of interest.

## SUPPLEMENTARY INFORMATION

The Supplementary Information associated with this paper includes Supplementary Synthetic Note, Supplementary Figures 1 – 13, Supplementary Tables 1 – 4.

