## Supplementary Information for "Chemical Proteomics Identifies Protein Ligands for Monoacylglycerol Lipids"

### TABLE OF CONTENTS


**Supplementary Figures 1 – 13** 3 – 15

**Supplementary Tables 1 – 4** 16 – 19

- See also separate excel sheets

**Supplementary References** 20

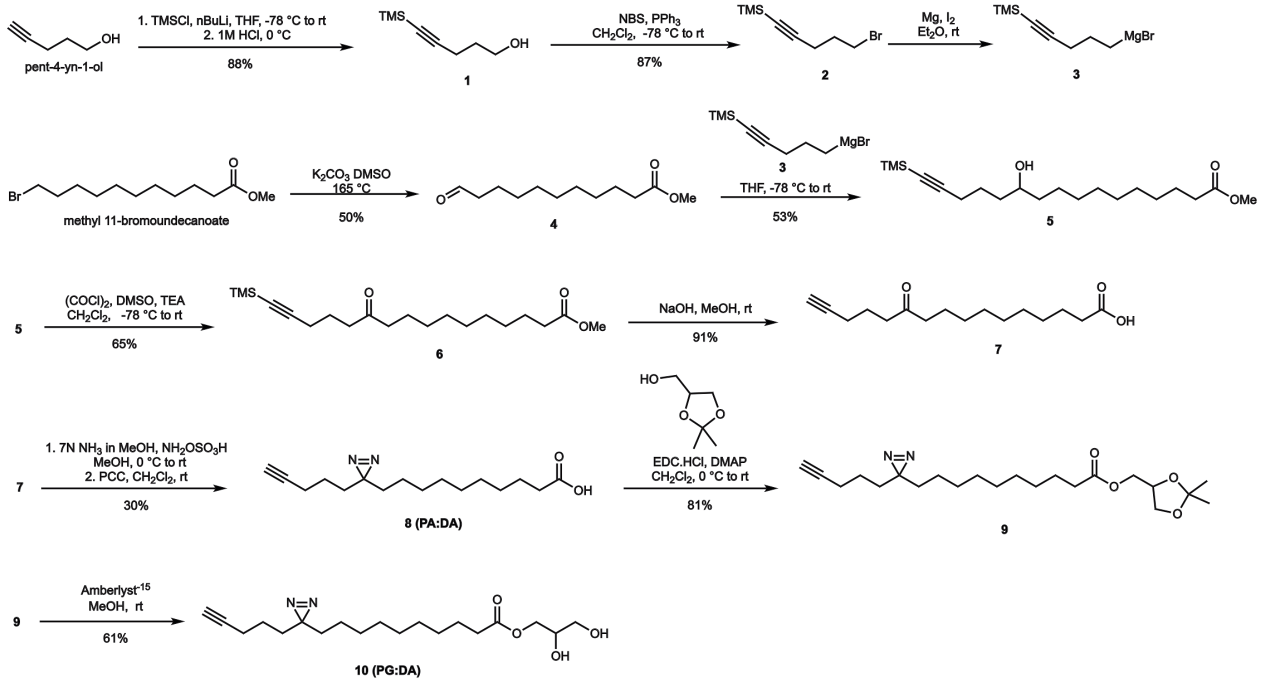


**Supplementary Figure 1.** The synthetic scheme used to generate the PG-DA probe. Complete details of all the synthesis and analytical characterization of the various intermediates can be found in the **Supplementary Synthetic Note**.

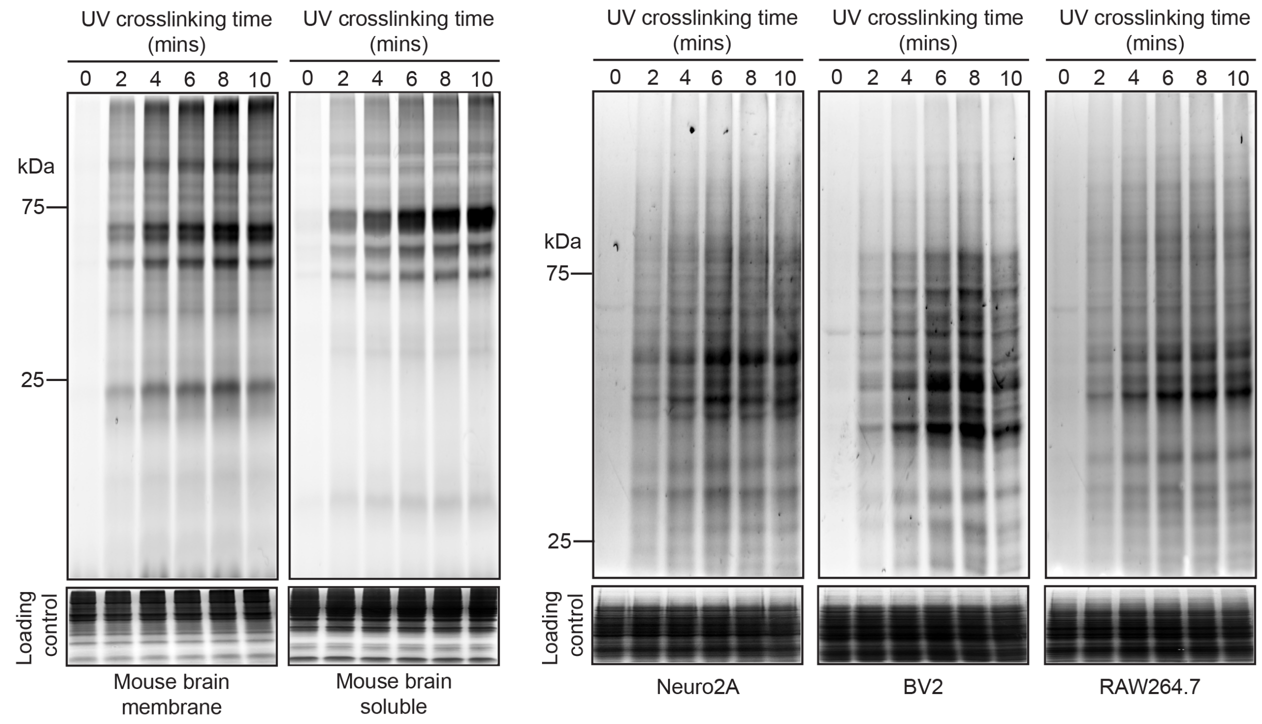


**Supplementary Figure 2.** UV-dependent crosslinking of the PG-DA probe (500 μM) in various lysates. In this experiment, UV-crosslinking time was varied from 0 – 10 mins, and in all cases, 6 mins was found to be the optimal time for UV crosslinking. The Coomassie staining shows the loading control for all gels in this experiment. This experiment was done three times with reproducible results each time.

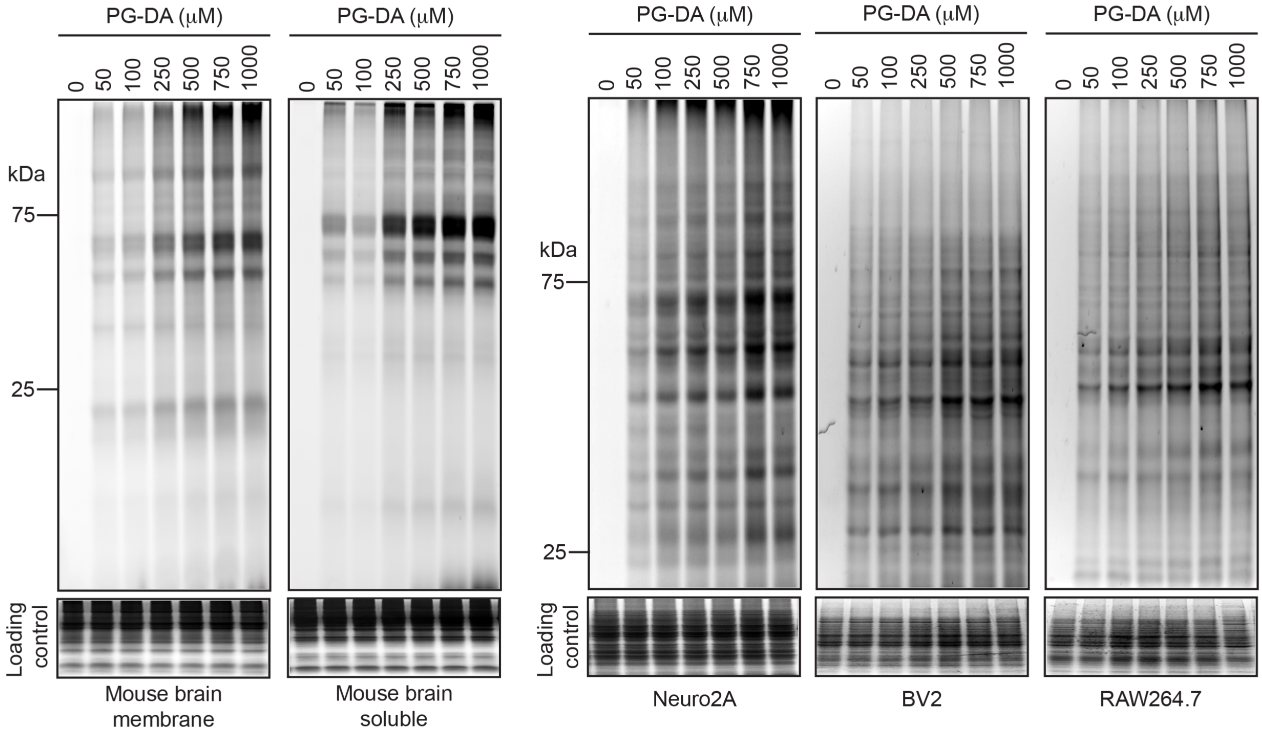


**Supplementary Figure 3.** UV-dependent crosslinking of various concentrations of the PG-DA probe (0 – 1000 μM) in various lysates. In this experiment, UV-crosslinking time was kept constant at 6 mins. The Coomassie staining shows the loading control for all gels in this experiment. This experiment was done three times with reproducible results each time.

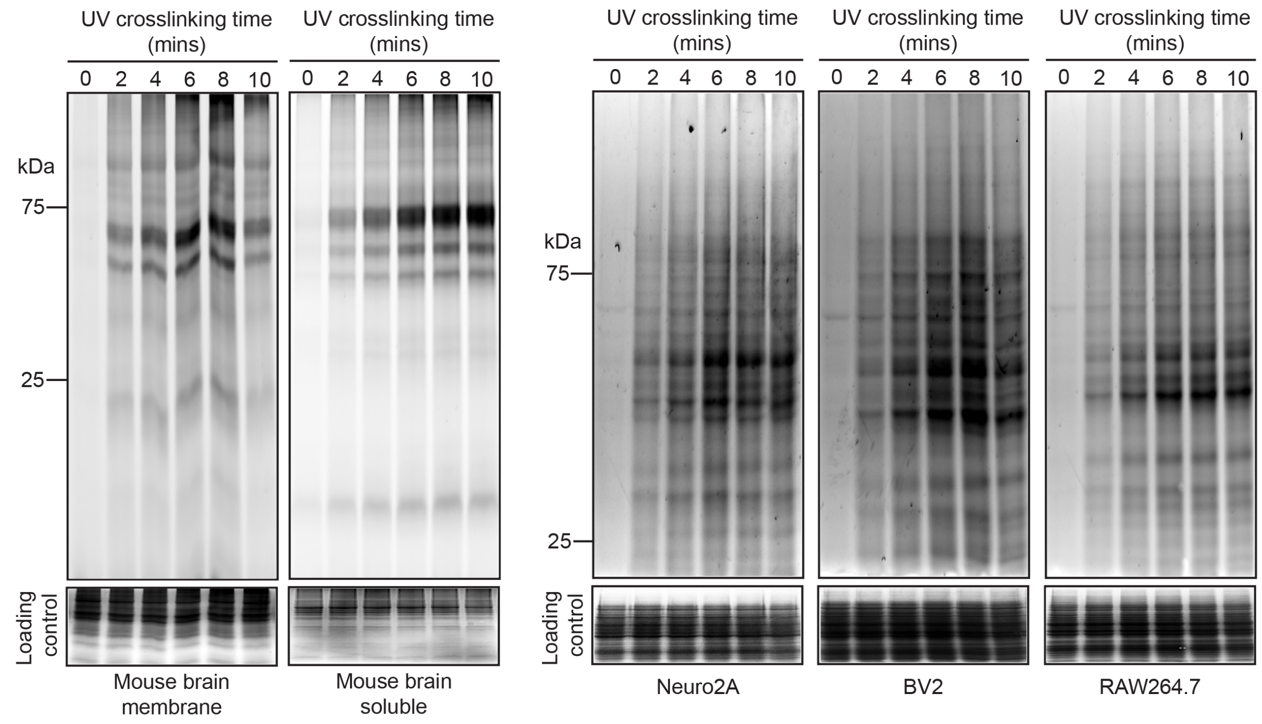


**Supplementary Figure 4.** UV-dependent crosslinking of the PA-DA probe (500 μM) in various lysates. In this experiment, UV-crosslinking time was varied from 0 – 10 mins, and in all cases, 6 mins was found to be the optimal time for UV crosslinking. The Coomassie staining shows the loading control for all gels in this experiment. This experiment was done three times with reproducible results each time.

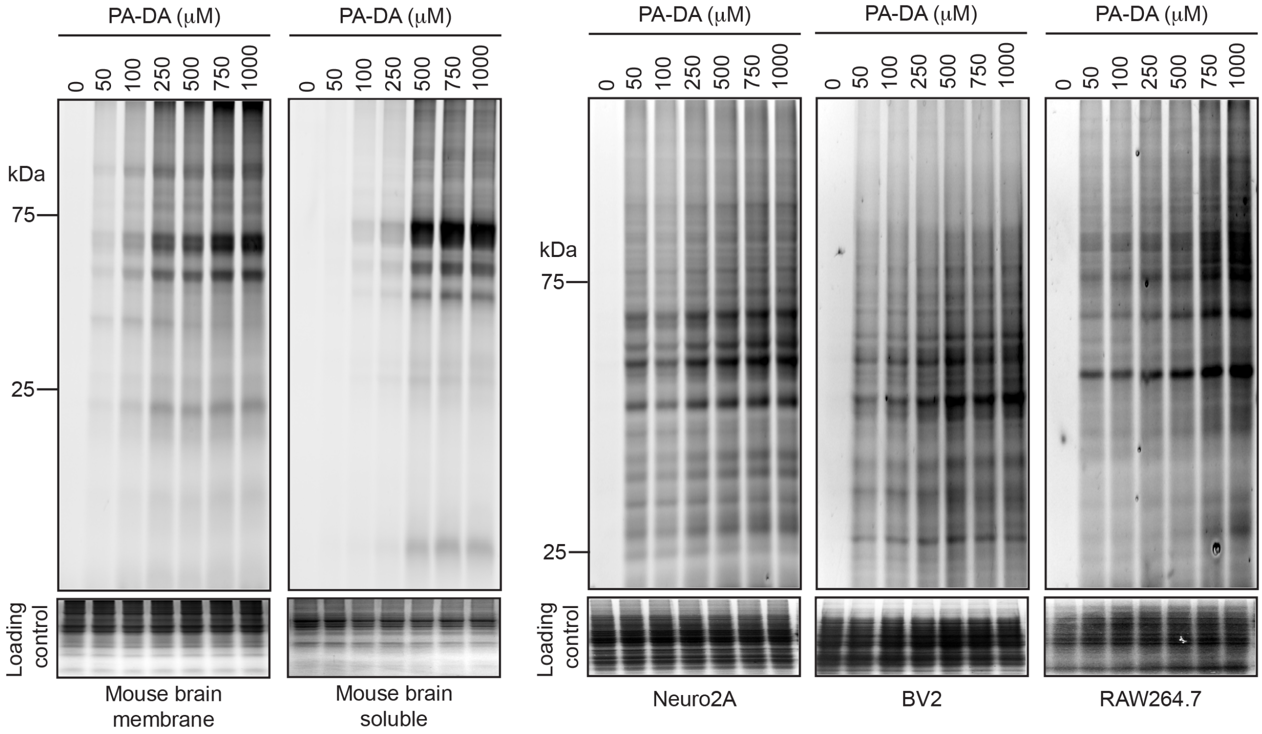


**Supplementary Figure 5.** UV-dependent crosslinking of various concentrations of the PA-DA probe (0 – 1000 μM) in various lysates. In this experiment, UV-crosslinking time was kept constant at 6 mins. The Coomassie staining shows the loading control for all gels in this experiment. This experiment was done three times with reproducible results each time.

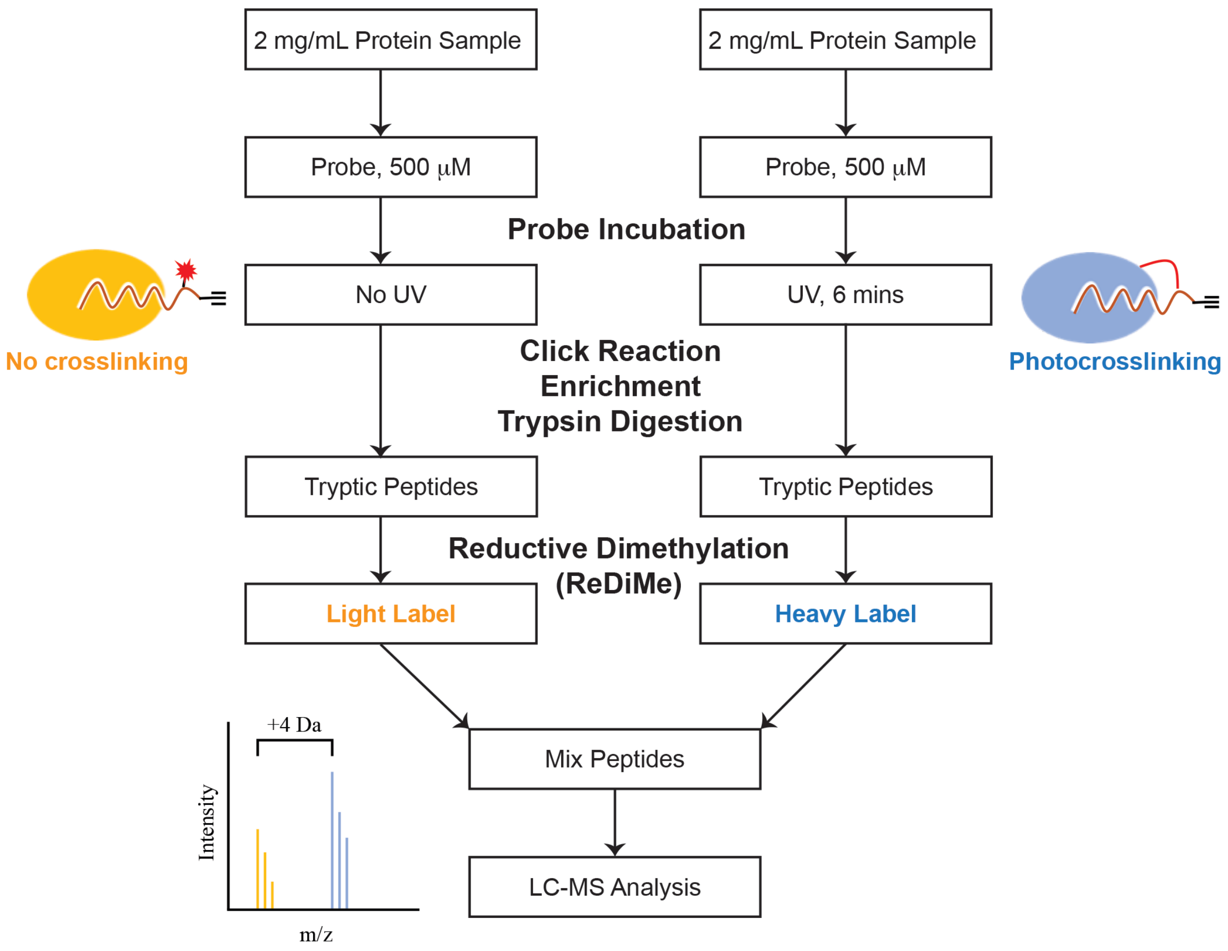


**Supplementary Figure 6.** A general workflow of the LC-MS/MS based quantitative chemoproteomics experiment for identifying total set of protein enriched by either the PG-DA or PA-DA probe in a UV-dependent manner from various mammalian lysates.

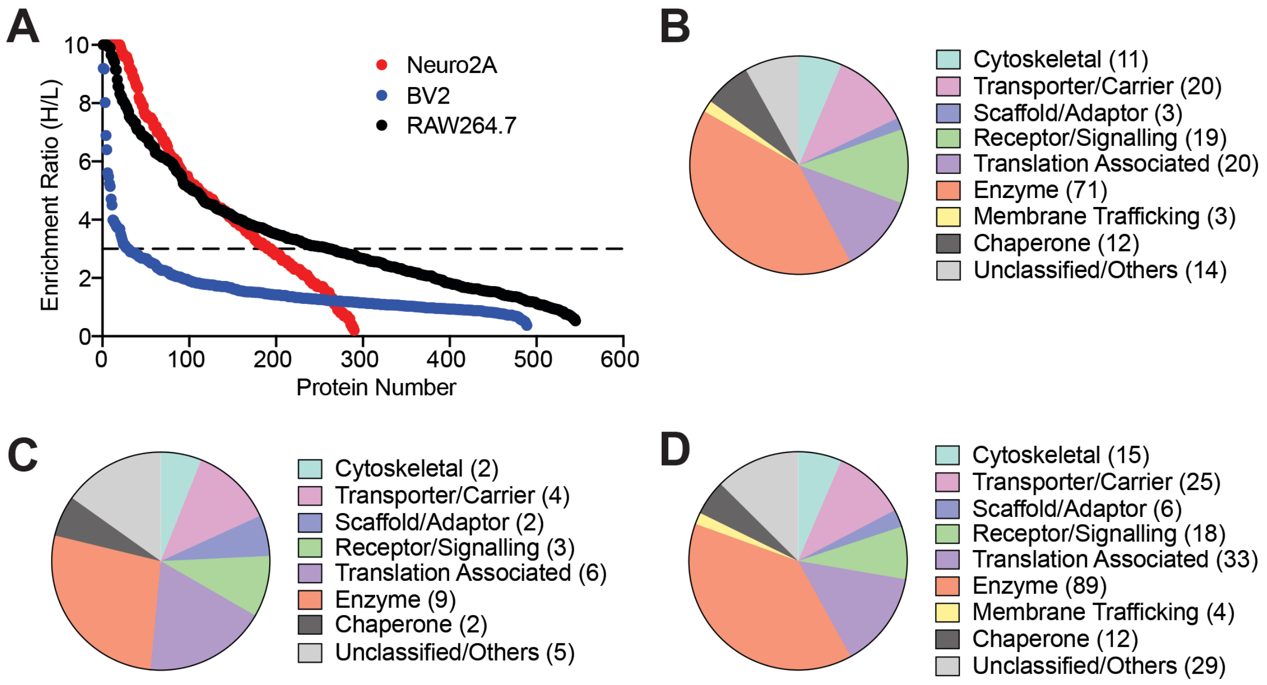


**Supplementary Figure 7.** (**A**) A LC-MS/MS based chemical proteomics experiment showing enrichment ratio (heavy:light; H:L) of the total proteins identified from the UV-dependent photocrosslinking of the PG-DA probe (500 μM, 6 mins of UV exposure) from the lysates of various immortalized mammalian cell lines. Each data point represents the mean of the enrichment ratio obtained for the respective protein from two or three biological replicate for a particular proteomic fraction, based on the defined filtering criteria for this proteomics experiment. The horizontal dotted line denotes an enrichment ratio ≥ 3, and proteins having an enrichment ratio above this threshold were considered enriched by the PG-DA probe, and taken forward for subsequent analysis. Complete details for all the proteins can be found in **Supplementary Table 1**. (**B–D**) Categorization of protein classes enriched by the PG-DA probe based on the Panther database annotation[1,2] for: (**B**) Neuro2A; (**C**) BV2; and (**D**) RAW264.7 cells respectively.

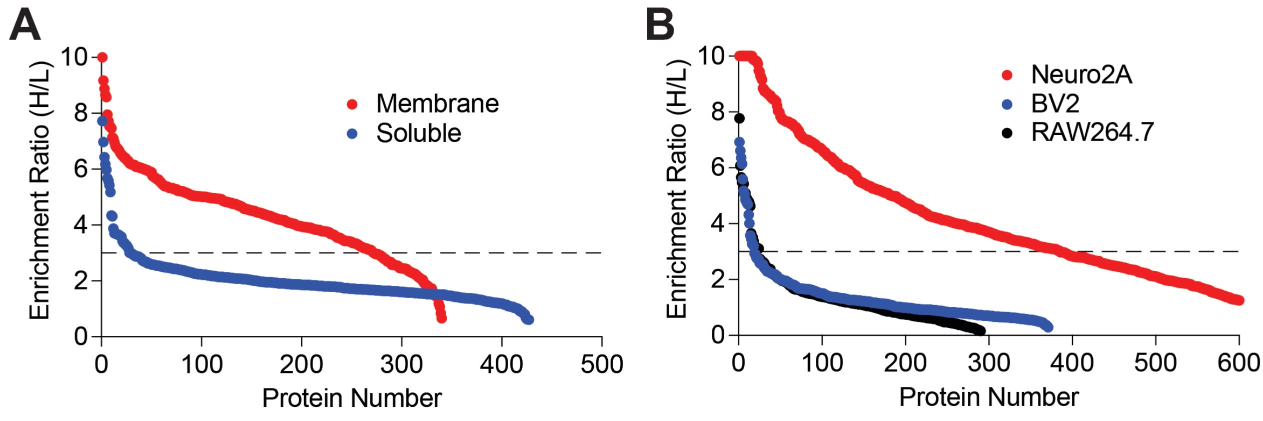


**Supplementary Figure 8.** A LC-MS/MS based chemical proteomics experiment showing enrichment ratio (heavy:light; H:L) of the total proteins identified from the UV-dependent photocrosslinking of the PA-DA probe (500 μM, 6 mins of UV exposure) from (**A**) the membrane and soluble lysates prepared from the mouse brain, and (**B**) from lysates of various immortalized mammalian cell lines. Each data point represents the mean of the enrichment ratio obtained for the respective protein from two or three biological replicate for a particular proteomic fraction, based on the defined filtering criteria for this proteomics experiment. The horizontal dotted line denotes an enrichment ratio ≥ 3, and proteins having an enrichment ratio above this threshold were considered enriched by the PA-DA probe, and taken forward for subsequent analysis. Complete details for all the proteins can be found in **Supplementary Table 2**.

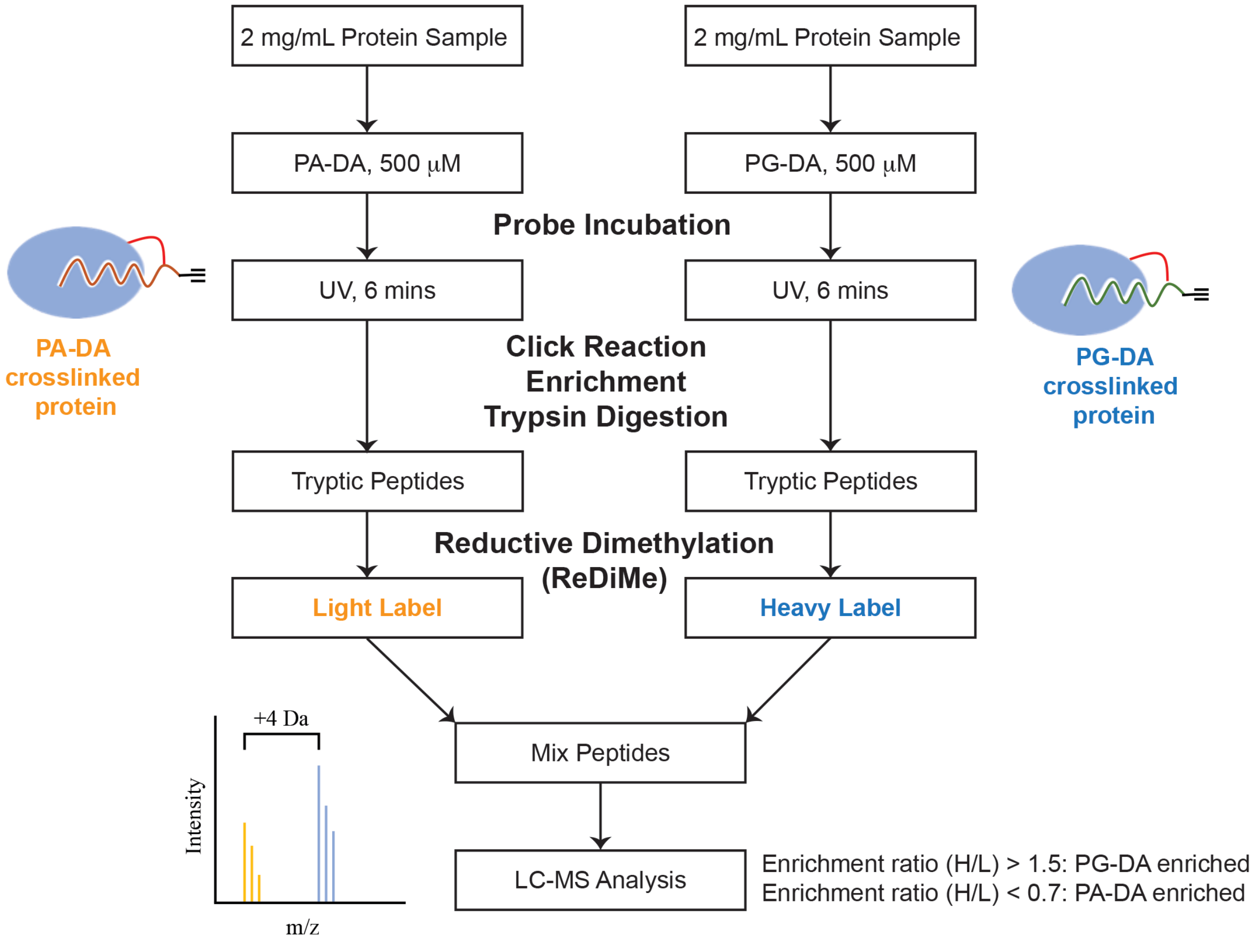


**Supplementary Figure 9.** A general workflow of the competitive LC-MS/MS based quantitative chemoproteomics experiment for identifying total set of protein enriched by either the PG-DA or PA-DA probe in a UV-dependent manner from various mammalian lysates.

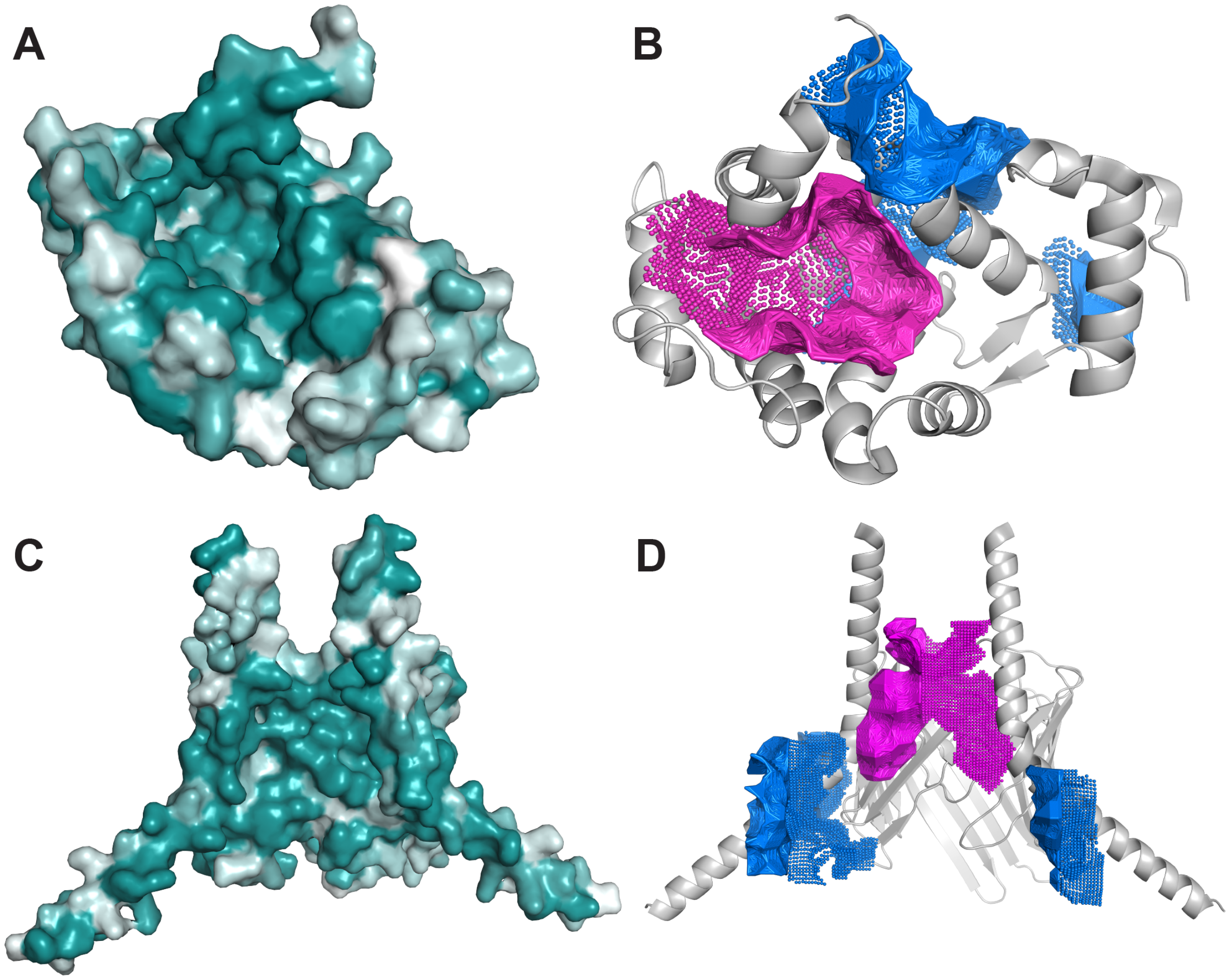


**Supplementary Figure 10.** Identification of potential hydrophobic 1-PG binding cavities in HPCA and TOMM22. (**A, B**) 1-PG binding cavities identified in HPCA using the (**A**) DEPTH[3], and (**B**) CavityPlus web servers[4]. (**C, D**) 1-PG binding cavities identified in HPCA using the (**C**) DEPTH[3], and (**D**) CavityPlus web servers[4].

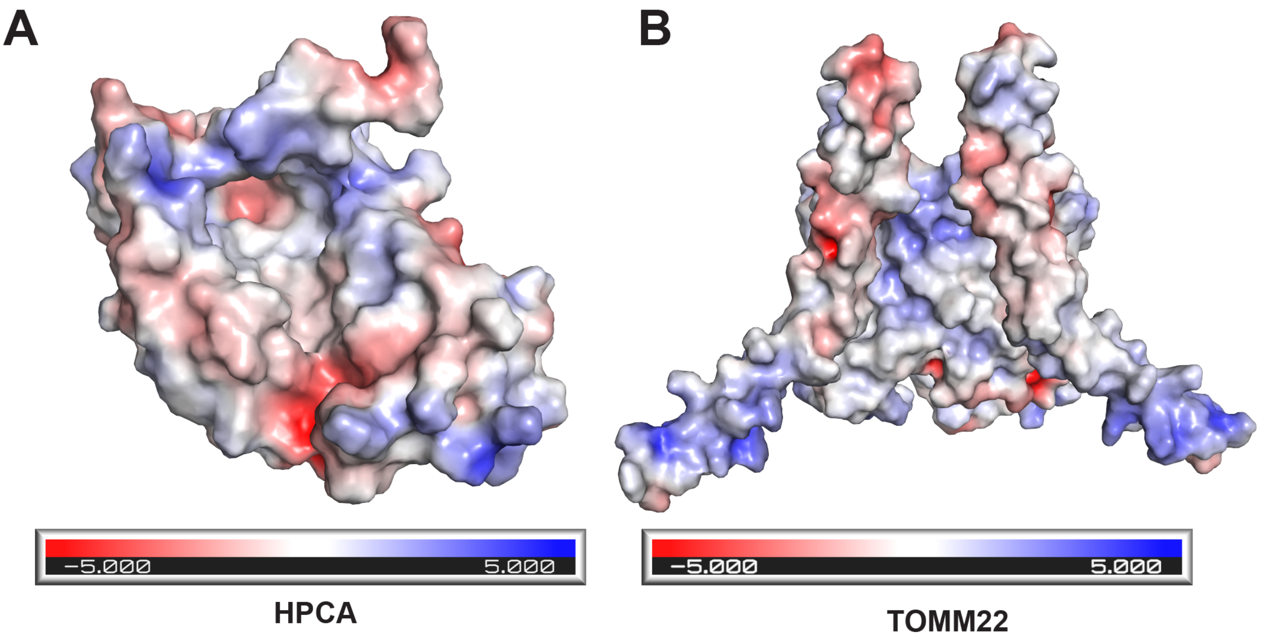


**Supplementary Figure 11.** Electrostatic surface map of (**A**) HPCA and (**B**) TOMM22, showing charges and hydrophobic pockets.

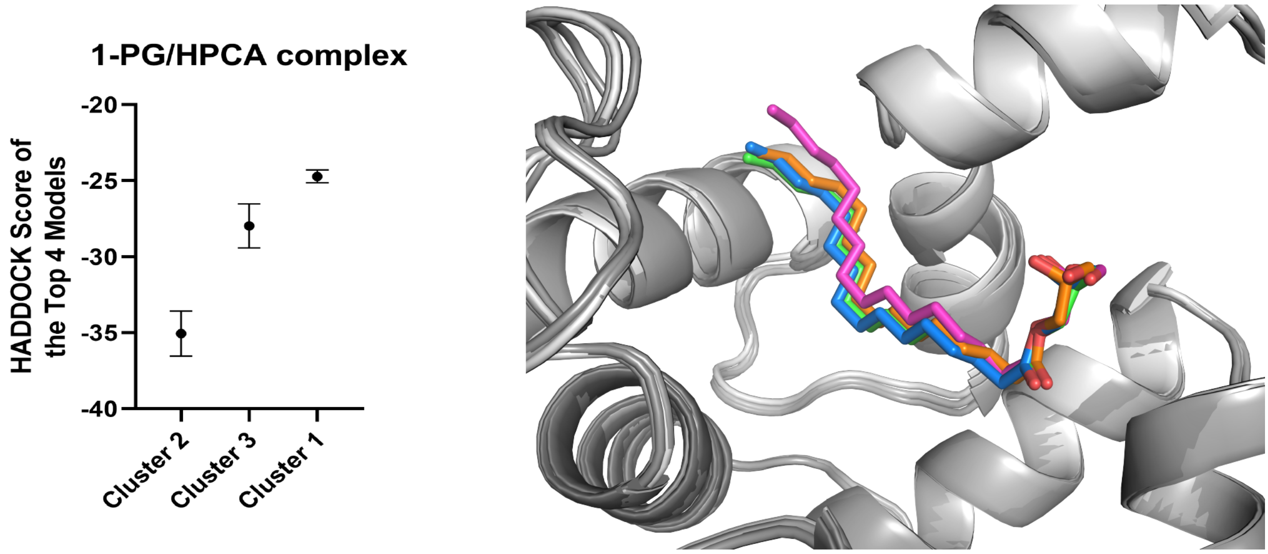


**Supplementary Figure 12.** (*Left*) The average HADDOCK score of the top four 1-PG/HPCA models in each cluster generate by HADDOCK[5]. (*Right*) Overlay of the top four 1-PG/HPCA models from the best cluster (cluster 2) showing ligand orientation in the binding pocket.

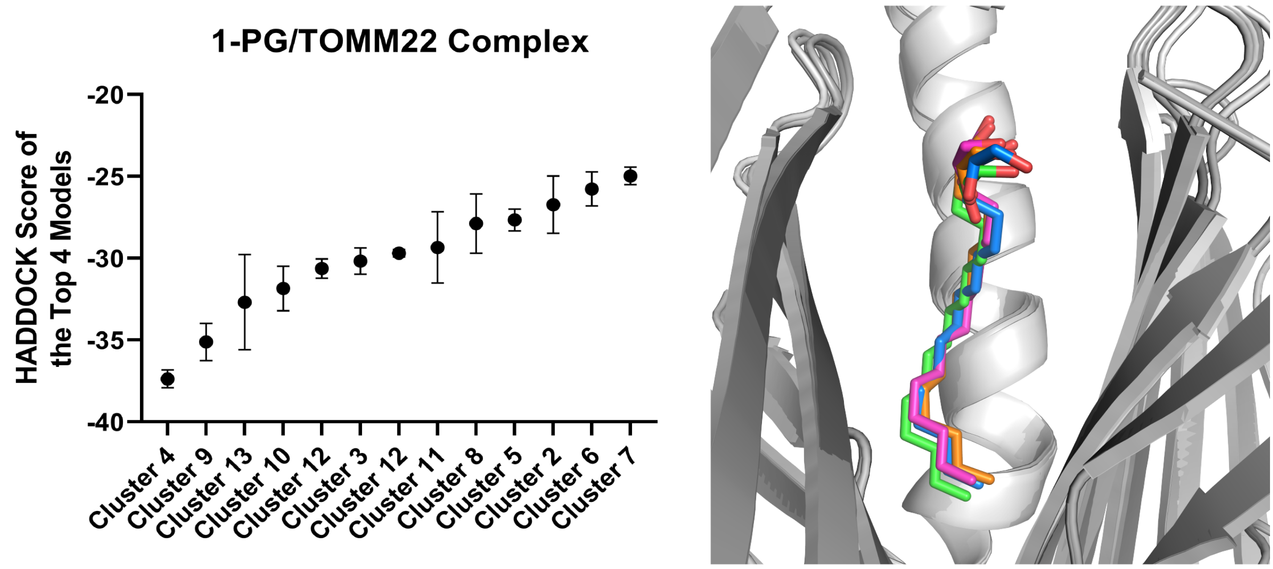


**Supplementary Figure 13.** (*Left*) The average HADDOCK score of the top four 1-PG/TOMM22 models in each cluster generate by HADDOCK[5]. (*Right*) Overlay of the top four 1-PG/TOMM22 models from the best cluster (cluster 9) showing ligand orientation in the binding pocket.

**SUPPLEMENTARY TABLE 1: A LC-MS/MS based chemoproteomic characterization of the PG-DA probe as a function of UV-crosslinking in different mammalian lysates.**

In all cases of these chemical proteomics experiments involving post-tryptic reductive demethylation (ReDiMe)[6], the tryptic peptides from the UV-crosslinked sample was labelled with heavy formaldehyde, while the tryptic peptides from the no UV sample was labelled with light formaldehyde. An enrichment ratio (H/L) represent relative abundance of a peptide in the heavy labelled sample versus the light labelled sample. For a protein to be classified as enriched by the PG-DA probe upon UV-crosslinking, it needed to be identified in at least 2 out of 3 replicates, have ≥ 3 quantified peptides per replicate, and have an enrichment ratio ≥ 3 in all replicates.

**Tab A**: List of proteins enriched by the PG-DA probe in the mouse brain membrane lysates.

**Tab B**: List of proteins enriched by the PG-DA probe in the mouse brain soluble lysates.

**Tab C**: List of proteins enriched by the PG-DA probe in the lysates obtained from Neuro2A cells.

**Tab D**: List of proteins enriched by the PG-DA probe in the lysates obtained from RAW264.7 cells.

**Tab E**: List of proteins enriched by the PG-DA probe in the lysates obtained from BV2 cells.

**SUPPLEMENTARY TABLE 2: A LC-MS/MS based chemoproteomic characterization of the PA-DA probe as a function of UV-crosslinking in different mammalian lysates.**

In all cases of these chemical proteomics experiments involving post-tryptic reductive demethylation (ReDiMe)[6], the tryptic peptides from the UV-crosslinked sample was labelled with heavy formaldehyde, while the tryptic peptides from the no UV sample was labelled with light formaldehyde. An enrichment ratio (H/L) represent relative abundance of a peptide in the heavy labelled sample versus the light labelled sample. For a protein to be classified as enriched by the PA-DA probe upon UV-crosslinking, it needed to be identified in at least 2 out of 3 replicates, have ≥ 3 quantified peptides per replicate, and have an enrichment ratio ≥ 3 in all replicates.

**Tab A**: List of proteins enriched by the PA-DA probe in the mouse brain membrane lysates.

**Tab B**: List of proteins enriched by the PA-DA probe in the mouse brain soluble lysates.

**Tab C**: List of proteins enriched by the PA-DA probe in the lysates obtained from Neuro2A cells.

**Tab D**: List of proteins enriched by the PA-DA probe in the lysates obtained from RAW264.7 cells.

**Tab E**: List of proteins enriched by the PA-DA probe in the lysates obtained from BV2 cells.

**SUPPLEMENTARY TABLE 3: A competitive LC-MS/MS based chemoproteomics experiments comparing the protein ligands of the PG-DA probe versus the PA-DA probe in different mammalian lysates.**

In all cases of these chemical proteomics experiments involving post-tryptic reductive demethylation (ReDiMe)[6], the tryptic peptides from the PG-DA treated sample was labelled with heavy formaldehyde, while the tryptic peptides from the PA-DA treated sample was labelled with light formaldehyde. An enrichment ratio (H/L) represent relative abundance of a peptide in the heavy labelled sample versus the light labelled sample. For a protein to be considered for any analysis in this experiment, it needed to be identified in at least 2 out of 3 replicates, and have ≥ 3 quantified peptides per replicate. A protein was considered enriched by the PG-DA probe, if it had an enrichment ratio ≥ 1.5 in all the replicates it was identified, while an enrichment ratio ≤ 0.7 classified a protein to be enriched by the PA-DA probe.

**Tab A**: Complete list of proteins identified in this competitive probe versus probe (PG-DA vs PA-DA) chemical proteomics experiments performed in the mouse brain membrane lysates.

**Tab B**: Complete list of proteins identified in this competitive probe versus probe (PG-DA vs PA-DA) chemical proteomics experiments performed in the mouse brain soluble lysates.

**Tab C**: Complete list of proteins identified in this competitive probe versus probe (PG-DA vs PA-DA) chemical proteomics experiments performed in the lysates from Neuro2A cells.

**Tab D**: Complete list of proteins identified in this competitive probe versus probe (PG-DA vs PA-DA) chemical proteomics experiments performed in the lysates from BV2 cells.

**Tab E**: Complete list of proteins identified in this competitive probe versus probe (PG-DA vs PA-DA) chemical proteomics experiments performed in the lysates from RAW264.7 cells.

**SUPPLEMENTARY TABLE 4. Datasets from the molecular docking of 1-PG into HPCA and TOMM22.**

**Tab A.** Identification of the top 5 cavities by CavityPlus based on the DrugScore for HPCA and TOMM22.

**Tab B**. Interaction energies for the 1-PG/HPCA and 1-PG/TOMM22 complexes for the top 4 models within the selected clusters.

**SUPPLEMENTARY REFERENCES.**


1 Thomas, P. D. *et al.* PANTHER: Making genome-scale phylogenetics accessible to all. *Protein Science* **31**, 8-22, doi:10.1002/pro.4218 (2022).

2 Mi, H., Muruganujan, A. & Thomas, P. D. PANTHER in 2013: modeling the evolution of gene function, and other gene attributes, in the context of phylogenetic trees. *Nucleic Acids Res* **41**, D377-386, doi:10.1093/nar/gks1118 (2013).

3 Tan, K. P., Nguyen, T. B., Patel, S., Varadarajan, R. & Madhusudhan, M. S. Depth: a web server to compute depth, cavity sizes, detect potential small-molecule ligand-binding cavities and predict the pKa of ionizable residues in proteins. *Nucleic Acids Res* **41**, W314-321, doi:10.1093/nar/gkt503 (2013).

4 Xu, Y. *et al.* CavityPlus: a web server for protein cavity detection with pharmacophore modelling, allosteric site identification and covalent ligand binding ability prediction. *Nucleic Acids Res* **46**, W374-W379, doi:10.1093/nar/gky380 (2018).

5 Honorato, R. V. *et al.* The HADDOCK2.4 web server for integrative modeling of biomolecular complexes. *Nature protocols*, doi:10.1038/s41596-024-01011-0 (2024).

6 Kelkar, D. S. *et al.* A chemical-genetic screen identifies ABHD12 as an oxidized-phosphatidylserine lipase. *Nat Chem Biol* **15**, 169-178, doi:10.1038/s41589-018-0195-0 (2019).
