## Supplementary Synthetic Note for "Chemical Proteomics Identifies Protein Ligands for Monoacylglycerol Lipids"

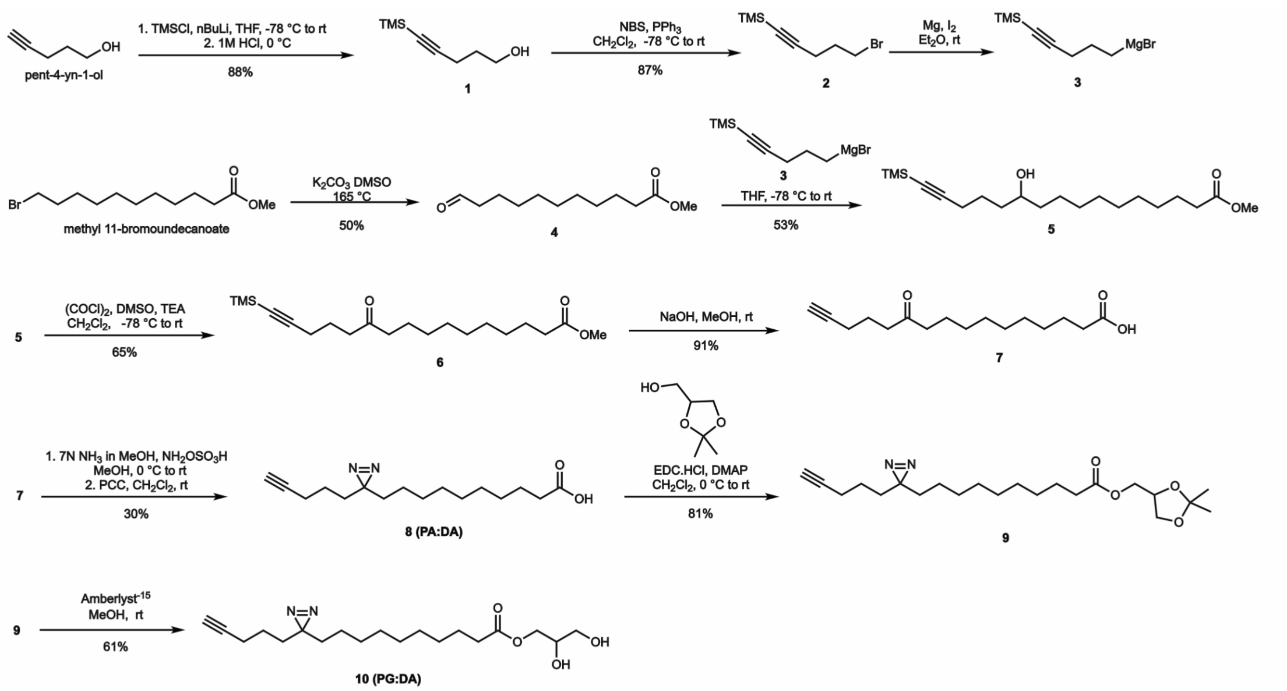


**Synthesis:** The bifunctional derivative, palmitoyl glycerol diazirine alkyne (PG-DA; compound **10**), features a palmitic acid chain with a terminal alkyne and an internal photoactive diazirine group (i.e., palmitic acid diazirine; PA-DA; compound **8**). A palmitic acid diazirine alkyne (**PA-DA**) was synthesized following previously established procedures with some modifications[1]. The reported synthesis of the aldehyde **4** used a lactone as a starting material, which is quite expensive and not readily commercially available. To address this shortcoming, we used a different approach. We started with an affordable and widely available alkyl halide, and converted it directly into the aldehyde using Kornblum oxidation[2]. Next, we encountered some difficulty with the earlier reported protocol[1] for the Grignard reaction using (5-chloropent-1-yn-1-yl)trimethylsilane. Specifically, a substantial portion of the compound remained unreacted while attempting to prepare the Grignard reagent from the TMS-protected pentynyl chloride. Considering the superior reactivity of a bromide compared to a chloride with magnesium, we decided to prepare (5-bromopent-1-yn-1-yl)trimethylsilane and used this as a substrate for the Grignard reaction. First, TMS protection of the alkyne of pent-4-yn-1-ol was carried out to give **1** in 88% yield. An Appel-type reaction[3] was used to convert the alcohol into the bromide **2** (87% yield). Treatment of a freshly prepared Grignard reagent **3** with aldehyde **4** afforded **5** in 53% yield. Subsequent steps were carried out using established protocols to produce **8** (**PA-DA**). For the synthesis of bifunctional MAG palmitic acid derivative, **PG-DA**, the following sequence of reactions was used. First, **8** (**PA-DA**) was treated with 1,2-isopropylideneglycerol in the presence of 1-ethyl-3-(3-dimethyl-aminopropyl)-carbodiimide hydrochloride (EDC.HCl) and the desired product **9** was isolated in 81% yield. Deprotection of the isopropylidene group of **9** using Amberlyst-15 gave **10** (**PG-DA**) in 69% yield.

**General.** All chemicals were purchased from Sigma-Aldrich and TCI, unless otherwise mentioned and used as received. All reactions were carried out under an atmosphere of nitrogen ($N_2$) or argon (Ar). Glassware was oven- or flame-dried prior to use. Analytical thin-layer chromatography (TLC) was performed using Silica Gel 60 F254 pre-coated plates (0.25 mm thickness, Merck), and visualized with staining with potassium permanganate ($KMnO_4$) or phosphomolybdic acid (PMA) solutions. Column chromatography was performed on Rankem silica gel (100-200 mesh). Nuclear magnetic resonance (NMR) spectra were recorded using deuterochloroform ($CDCl_3$), as the solvent. $^1H$, $^{13}C$ spectra were recorded on JEOL 400 MHz or Bruker 400 MHz (or 100 MHz for $^{13}C$) NMR spectrometers. The internal standards for the recorded NMR spectra were either residual solvent signals (Chloroform, $\delta_H$ = 7.26 ppm, $\delta_C$ = 77.2 ppm and or an internal standard tetramethylsilane ($\delta_H$ =0.00 ppm, $\delta_C$ = 0.00 ppm). Chemical shifts ($\delta$) are reported in ppm and coupling constants ($J$) in Hz, multiplicities were reported by the following abbreviations: s (singlet), broad singlet (bs), d (doublet), dd (doublet of doublet), t (triplet), q (quartet), m (multiplet). High-resolution mass spectra were obtained from high-resolution mass spectrometry (HRMS)–electrospray ionization–Q-TOF–LC–MS/MS (Sciex)

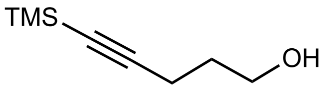


**5-(Trimethylsilyl)pent-4-yn-1-ol (1).** Compound **1** was synthesized according to a procedure reported previously (Harris et al. 2014) with some modifications[4], and the analytical data collected by us is consistent with literature values. Briefly, in a flame-dried, two-necked round-bottom flask, containing 4-pentyn-1-ol (5.0 g,59.44 mmol, 1.0 equiv.) and anhydrous THF (100 mL) were added, and the mixture was cooled to -78 ºC. To this, *n*-butyllithium (52 mL, 2.5 M in hexanes, 130.76 mmol, 2.2 equiv.) was added dropwise over a period of 30 min and the temperature was maintained at -78 ºC. Next, chlorotrimethylsilane (22.63 mL, 178.32 mmol, 3.0 equiv.) was added dropwise, and the reaction mixture was gradually warmed to room temperature and stirred for an additional 16 h. The reaction mixture was subsequently cooled to 0 ºC and acidified with 1 M HCl (150 mL), followed by stirring for 1 h. The resulting mixture was then extracted with $Et_2O$(3 x 100 mL), and the combined organic layer was washed with water (250 mL), sat. $NaHCO_3$ (200 mL), and brine (200 mL), dried over $Na_2SO_4$, and concentrated *in vacuo*. The obtained residue was purified by silica gel column chromatography (elution with gradient 10–25% EtOAc/ hexanes) to afford the compound **1** as a colourless oil (8.17 g, 88%). **TLC** $R_f$ = 0.2 (10% EtOAc/ Hexanes; TLC stain, $KMnO_4$); **$^1$H NMR** (400 MHz, $CDCl_3$): $\delta$ 3.75 (t, $J$ = 6.1 Hz, 2H), 2.34 (t, $J$ = 6.9 Hz, 2H), 1.76 (quint, $J$ = 6.6 Hz, 2H), 1.72 (bs, 1H), 0.14 (s, 9H); **HRMS** (*m/z*): [M + H]+ calcd**.** for $C_8H_{17}OSi$, 157.1048; found, 157.1051.

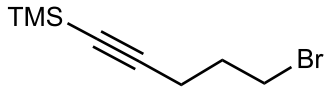

**(5-Bromopent-1-yn-1-yl)trimethylsilane (2).** Compound **2** was synthesized according to a procedure reported previously (Yao et al. 2002) with some modifications[5], and the analytical data collected by us is consistent with literature values. To a solution of compound **1** (5.1 g, 32.63 mmol, 1 equiv.) in anhydrous $CH_2Cl_2$ (100 mL). Triphenylphosphine (10.27 g, 39.16 mmol, 1.2 equiv.) was added portion wise at –78 °C under $N_2$ atmosphere. *N*-Bromosuccinimide (6.39 g, 35.89 mmol, 1.1 equiv.) was then slowly added, and the reaction mixture was gradually warmed to room temperature and stirred for 6 h. After completion of reaction (TLC analysis), $Et_2O$ (400 mL) was added to the reaction mixture and washed with sat. $NaHCO_3$ (2 x 200 mL), brine (200 mL), dried over $Na_2SO_4$, and concentrated *in vacuo*. The resulting crude was purified by silica gel column chromatography (elution with 100% hexanes) to afford the **2** as a colourless oil (6.22 g, 87%). **TLC** $R_f$ = 0.5 (100% hexanes; TLC stain, $KMnO_4$) **$^1$H NMR** (400 MHz, $CDCl_3$): $\delta$ 3.51 (t, $J$ = 6.5 Hz, 2H), 2.41 (t, $J$ = 6.8 Hz, 2H), 2.04 (quint, $J$ = 6.6 Hz, 2H), 0.15 (s, 9H).

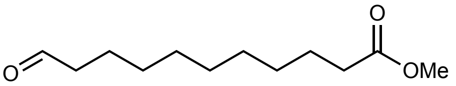

**Methyl 11-oxoundecanoate (4).** Compound **4** was synthesized according to a procedure reported previously (Ravi, S. et al. 2001) with some modifications[2], and the analytical data collected by us is consistent with literature values. To a solution of methyl 11-bromoundecanoate (0.5 g, 1.791 mmol, 1 equiv.) in anhydrous DMSO (5 mL), $NaHCO_3$ (0.33 g, 3.94 mmol, 2.2 equiv.) was added. The reaction mixture was refluxed at 165 °C for 15 min and then cooled to room temperature, diluted with ice cold water (10 mL), and extracted with $Et_2O$ (2 x 25 mL). The combined organic layer was washed with water (25 mL), sat. $NaHCO_3$ (20 mL), and brine (20 mL). The resulting organic layer was dried over $Na_2SO_4$, and concentrated *in vacuo*. The crude was purified by silica gel column chromatography (elution with gradient 10–25% EtOAc/ hexanes) to afford the compound **4** as a colourless oil (0.19 g, 50%). **TLC** $R_f$ = 0.5 (10% EtOAc/ hexane; TLC stain, $KMnO_4$); **$^1$H NMR** (400 MHz, $CDCl_3$): $\delta$ 9.76 (t, $J$ = 1.9 Hz, 1H), 3.66 (s, 3H), 2.41(td, $J$ = 77.4, 1.9 Hz, 2H), 2.30 (t, $J$ = 7.5 Hz, 2H), 1.66–1.58 (m, 4H), 1.36–1.25 (m, 10H); **HRMS** (*m/z*): [M + H]+ calcd. for $C_{12}H_{23}O_3$, 215.1647; found, 215.1636.

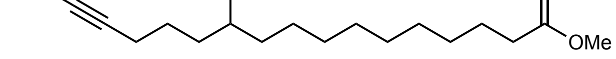

**Methyl 11-hydroxy-16-(trimethylsilyl)hexadec-15-ynoate (5).** <u>*Preparation of Grignard reagent* (**3**)</u>*:* The Grignard reagent was prepared based on the procedure reported by (Harris et al. 2014) with modifications[4]. Reaction flasks, glass syringes, needles and magnetic stir bars were dried at 110 °C in an oven for 2 h. Magnesium turnings were treated with a 10% aq. HCl solution, followed by subsequent rinsing with water and acetone. The magnesium turnings were then dried in the oven at 110 ºC for 2 h. Before commencement of the reaction, all items in the oven were gradually cooled to room temperature in a desiccator. The solvents used for the reaction $Et_2O$ and tetrahydrofuran (THF) were dried over sodium metal and then freshly distilled. Also freshly distilled bromide **2** (5-10 mm Hg (Torr), 85-90 ºC; 70-80% yield) was used for the preparation for the Grignard reagent. A 100 mL Schlenk round-bottom flask was charged with the magnesium turnings (0.35 g, 14.45 mmol, 1.2 equiv.) and heated with a heat gun for 10 min under vacuum, and then cooled to room temperature under $N_2$ atmosphere. A magnetic stir bar was introduced into the flask, and anhydrous $Et_2O$ (16 mL) was added, followed by the addition of a catalytic amount of iodine (8.6 mg, 0.068 mmol, 0.006 equiv.). The mixture was stirred at room temperature for 10 min. A solution of **2** (2.64 g, 12.04 mmol, 1.0 equiv.) in $Et_2O$ (16 mL) was prepared and a small portion, approximately 20%, was added dropwise, and then reaction mixture was stirred at room temperature for 10-15 min until the colour of the solution transitioned from brown/red to colorless, signifying the initiation of the Grignard reagent. The remaining solution of **2** was then added dropwise over 15 min and the reaction mixture was stirred at room temperature for another 3 h. The Grignard reagent **3** was used immediately for the next step.

_Grignard reaction:_ A solution of methyl 11-oxoundecanoate **4** (2.2 g, 10.27 mmol, 1.0 equiv.) in anhydrous THF (33 mL) was cooled to -78 ºC under $N_2$ atmosphere. The Grignard solution **3** was added dropwise to the stirring solution over 15 min. The reaction mixture was then gently warmed to 0 ºC and stirred for 1 h. Saturated $NH_4Cl$ solution (250 mL) was used to quench the reaction. The product was extracted with EtOAc (3 x 100 mL), and the combined organic layers was dried with $Na_2SO_4$, and concentrated in *vacuo*. The resulting crude was purified by silica gel column chromatography (elution with gradient 10–20% EtOAc/ hexanes) to afford the compound **5** as a colorless oil (1.93 g, 53%). **TLC** $R_f$ = 0.4 (20% EtOAc/ hexanes, stain with $KMnO_4$); The analytical data collected by us is consistent with reported literature (Hulce et al. 2013)[1] values: **$^1$H NMR** (400 MHz, $CDCl_3$): $\delta$ 3.64 (s, 3H), 3.63–3.55 (m, 1H), 2.28 (t, *J* = 7.5 Hz, 2H), 2.23 (t, *J* = 6.5 Hz, 2H), 1.63–1.38 (m, 8H), 1.27 (d, *J* = 4.6 Hz, 12H), 0.12 (s, 9H); **HRMS** (*m/z*): [M + H]+ calcd. for $C_{20}H_{39}O_3Si$, 355.2668; found, 355.2667.

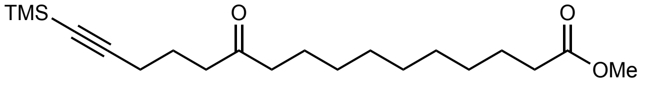


**Methyl 11-oxo-16-(trimethylsilyl)hexadec-15-ynoate (6).** Compound **6** was synthesized according to a procedure reported previously (Hulce et al. 2013)[1] with some modifications, and the analytical data collected by us is consistent with literature values. _Swern Oxidation:_ To a stirring solution of Oxalyl chloride (0.87 mL, 10.16 mmol, 2.0 equiv.) in anhydrous $CH_2Cl_2$ (65 mL), DMSO (1.45 mL, 20.32 mmol, 4.0 equiv.) was added dropwise at -78 ºC and the reaction mixture was stirred for 30 min. Subsequently, solution of compound **5** (1.8 g, 5.08 mmol, 1.0 equiv) in $CH_2Cl_2$ (5 mL) was added dropwise, and the reaction mixture was stirred for another 30 min at -78 ºC. Lastly, triethylamine (2.84 mL, 20.32 mmol, 4 equiv.) was added, and after 30 min, the reaction mixture was gently raised to 0 ºC and stirred at this temperature for 1 h. The reaction mixture was then diluted with $H_2O$ (300 mL) and extracted in $CH_2Cl_2$ (3 x 100 mL). The combined organic layer was washed with brine (150 mL) and dried over $Na_2SO_4$ and concentrated in *vacuo*. The crude residue was purified by silica gel column chromatography (elution with gradient 5–10% EtOAc/ hexanes) to afford the compound **6** as a white solid (1.17 g, 65% yield): **TLC** $R_f$ = 0.4 (10% EtOAc/ hexanes; TLC stain, $KMnO_4$); **$^1$H NMR** (400 MHz, $CDCl_3$): $\delta$ 3.64 (s, 3H), 2.53 (t, *J* = 7.2 Hz, 2H), 2.40 (t, *J* = 7.4 Hz, 2H), 2.28 (t, *J* = 7.5 Hz, 2H), 2.24 (t, *J* = 6.9 Hz, 2H), 1.75 (quintet, *J* = 7.0 Hz, 2H), 1.64–1.52 (m, 4H), 1.32–1.22 (m, 10H), 0.13 (s, 9H); **HRMS** (*m/z*): [M + H]+ calcd. for $C_{20}H_{37}O_3Si$ [M + H]+ 353.2512, found, 353.2507.

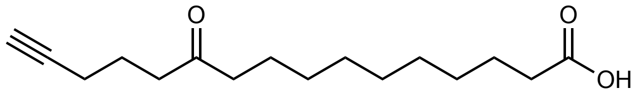


**11-Oxohexadec-15-ynoic acid (7).** Compound **7** was synthesized according to a procedure reported previously (Hulce et al. 2013) with some modifications[1], and the analytical data collected by us is consistent with literature values. To a solution of **6** (1.16 g, 3.29 mmol, 1.0 equiv.) in MeOH (10 mL), NaOH (0.66 g, 16.45 mmol, 5 equiv.) in $H_2O$ (10 mL) was added dropwise, and the reaction mixture was stirred at room temperature for 16 h. The reaction mixture was acidified to pH 2-3 by treating with 10% aq. HCl at 0 ºC, and extracted with $Et_2O$ (3 x 200 mL). The combined organic layer was washed with brine (250 mL) and dried over $Na_2SO_4$, and concentrated in vacuo The crude was purified by silica gel column chromatography (elution with gradient 15–25% EtOAc/hexanes, 1% $HCO_2H$) to afford the compound **7** as a white solid (798 mg, 91%). **TLC** $R_f$ = 0.2 (20% EtOAc/ hexanes with 1% $HCO_2H$; TLC stain, PMA); **$^1$H NMR** (400 MHz, $CDCl_3$): $\delta$ 2.55 (t, *J* = 7.2 Hz, 2H), 2.40 (t, *J* = 7.4 Hz, 2H), 2.34 (t, *J* = 7.5 Hz, 2H), 2.22 (td, *J* = 6.9, 2.8 Hz, 2H), 1.95 (t, *J* = 2.6 Hz, 1H), 1.78 (quintet, *J* = 7.1 Hz, 2H), 1.66–1.52 (m, 4H), 1.34–1.23 (m, 10H); **HRMS** (*m/z*): [M + Na]+ *m/z* calcd. for $C_{16}H_{26}O_3Na$, 289.1780; found 289.1788.

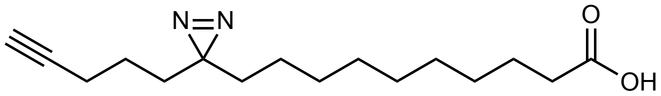


**10-(3-(Pent-4-yn-1-yl)-3H-diazirin-3-yl)decanoic acid (PA-DA) (8)** Compound **8** (**PA-DA**) was synthesized according to a procedure reported previously (Hulce et al. 2013) with some modifications[1], and the analytical data collected by us is consistent with literature values. The reaction and column chromatography were conducted in the dark due to the light sensitivity of the product. Briefly, in a sealed tube containing a stir bar, 11-oxohexadec-15-ynoic acid (200 mg, 0.75 mmol, 1.0 eq.) was added and cooled to 0 ºC. 7 N $NH_3$ in MeOH (5.8 mL) was added dropwise and the resulting reaction mixture was stirred for 3 h, while maintaining the temperature at 0 ºC. A solution of hydroxylamine-*O*-sulfonic acid (97 mg, 0.86 mmol, 1.15 equiv.) in MeOH (2.5mL) was gradually added dropwise to the reaction mixture. The seal tube was covered with aluminium foil and the reaction mixture was stirred at room temperature overnight, and then the solvent was evaporated under a stream of nitrogen gas ($N_2$). To the remaining residue, $Et_2O$ (10.0 mL) was added, leading to the formation of a suspension containing insoluble salts, which were subsequently filtered off. The filtrate ($Et_2O$ layer) was concentrated under reduced pressure, and the residue obtained was re-dissolved in anhydrous $CH_2Cl_2$ (12.0 mL) and pyridine (1.75 mL). To this mixture, pyridinium chlorochromate (PCC, 485 mg, 2.25 mmol, 3.0 equiv.) was then added, and the reaction mixture was stirred for 3 h at room temperature. Subsequently, the reaction mixture was passed through a silica pad using a solvent mixture of 70% EtOAc/ hexanes containing 1% formic acid ($HCO_2H$), and the resulting solution was concentrated *in vacuo*, and the crude was purified by silica gel column chromatography (elution with gradient 10–15% EtOAc/hexanes, 1% $HCO_2H$) to afford the compound **8** as a white solid (63 mg, 30%). **TLC** $R_f$ = 0.4 (30% EtOAc/hexanes with 1% $HCO_2H$; TLC stain PMA] **$^1$H NMR** (400 MHz, $CDCl_3$): $\delta$ 2.33 (t, $J$ = 7.5 Hz, 2H), 2.15 (td, $J$ = 6.9, 2.6 Hz, 2H), 1.94 (t, $J$ = 2.6 Hz, 1H), 1.61 (quintet, $J$ = 7.5 Hz, 2H), 1.52–1.43 (m, 2H), 1.39–1.15 (m, 14H), 1.12–1.01 (m, 2H); **HRMS** (*m/z*): [M + H]+ calcd. for $C_{16}H_{27}N_2O_2$, 279.2073; found, 279.2071.

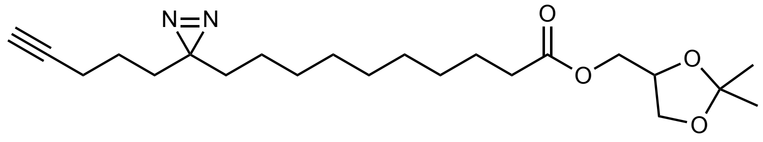


**2,2-Dimethyl-1,3-dioxolan-4-yl)methyl 10-(3-(pent-4-yn-1-yl)-3H-diazirin-3-yl)decanoate (9)** To a solution of **8** (**PA-DA**, 48 mg, 0.17 mmol, 1.0 equiv.) and 1,2-isopropylideneglycerol (23 mg, 0.17 mmol, 1.0 equiv.) in anhydrous $CH_2Cl_2$ (4.8 mL), 4-dimethylaminopyridine (5 mg, 0.04 mmol, 0.25 equiv.) was added. The mixture was cooled to 0 ºC and EDC.HCl (0.05 g, 0.26 mmol, 1.5 equiv.) was added in a single portion. The reaction mixture was gradually warmed to room temperature and stirred for 16 h. After completion of reaction (TLC analysis), the reaction was diluted with $CH_2Cl_2$ (50 mL) and washed with sat. $NaHCO_3$ (2 x 20 mL) and brine (50 mL). The combined organic layer was dried over $Na_2SO_4$, and concentrated *in vacuo*. The crude residue was purified by silica gel column chromatography (elution with gradient 5–10% EtOAc/ hexanes) to afford **9** as a colorless oil (55 mg, 81%). **TLC** $R_f$ = 0.3 (10% EtOAc/ hexanes; TLC stain, PMA); **$^1$H NMR** (400 MHz, $CDCl_3$): $\delta$ 4.35–4.27 (m, 1H), 4.16 (dd, $J$ = 11.5, 4.7 Hz, 1H), 4.12–4.04 (m, 2H), 3.73 (dd, $J$ = 8.4, 6.2 Hz, 1H), 2.33 (t, $J$ = 7.5 Hz, 2H), 2.16 (td, $J$ = 6.9, 2.6 Hz, 2H), 1.94 (t, $J$ = 2.7 Hz, 1H), 1.65–1.56 (m, 2H), 1.51–1.45 (m, 2H), 1.43 (s, 3H), 1.38–1.18 (m, 17H), 1.12–1.01 (m, 2H); **$^{13}$C NMR** (100 MHz, $CDCl_3$): $\delta$ 173.8, 110.0, 83.6, 73.8, 69.0, 66.5, 64.7, 34.2, 33.0, 32.0, 29.4, 29.4, 29.3, 29.3, 29.2, 28.6, 26.8, 25.5, 25.0, 23.9, 22.9, 18.1; **HRMS** (*m/z*): [M + H]+ calcd. for $C_{22}H_{37}N_2O_4$, 393.2753; found, 393.2752.

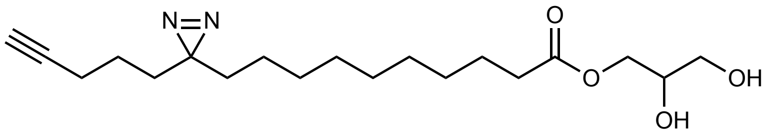

**2,3-Dihydroxypropyl 10-(3-(pent-4-yn-1-yl)-3H-diazirin-3-yl)decanoate (PG-DA) (10).** Amberlyst-15 ($H^+$ form, 88 mg, 0.28 mmol, 2 equiv.) was added to a solution of **compound 9** (55 mg, 0.140 mmol, 1 equiv.) in MeOH (9.0 mL). The resulting reaction mixture was stirred for 18 h at room temperature. After completion of reaction (TLC analysis), Amberlyst-15 was filtered off, and the filtrate was concentrated *in vacuo*. The crude residue was purified by silica gel column chromatography (elution with gradient 30–60% EtOAc/ hexanes) to afford the desired compound **10 (PG-DA)** as a colorless oil (30 mg, 61%). **TLC** $R_f$ = 0.5 (60% EtOAc/hexanes; TLC stain, PMA); **$^1$H NMR** (400 MHz, $CDCl_3$): $\delta$ 4.21–4.07 (m, 2H), 3.95–3.87 (m, 1H), 3.68 (dd, *J* = 11.5, 3.9 Hz, 1H), 3.58 (dd, *J* = 11.5, 5.9 Hz, 1H), 2.91 (bs, 1H), 2.56 (bs, 1H), 2.33 (t, *J* = 7.6 Hz, 2H), 2.14 (td, *J* = 7.0, 2.6 Hz, 2H), 1.94 (t, *J* = 2.7 Hz, 1H), 1.67–1.54 (m, 2H), 1.51–1.42 (m, 2H), 1.38–1.15 (m, 14H), 1.11–1.00 (m, 2H).;**$^{13}$C NMR** (100 MHz, $CDCl_3$) $\delta$ 174.5, 83.6, 70.4, 69.0, 65.2, 63.5, 34.2, 32.9, 31.9, 29.4, 29.3, 29.2, 29.2, 29.2, 28.6, 25.0, 23.9, 22.9, 18.1; **HRMS** *(m/z):* [M + H]+ calcd. for $C_{19}H_{33}N_2O_4$, 353.2440; found 353.2437.

**¹H NMR of 5-(Trimethylsilyl)pent-4-yn-1-ol (1)**

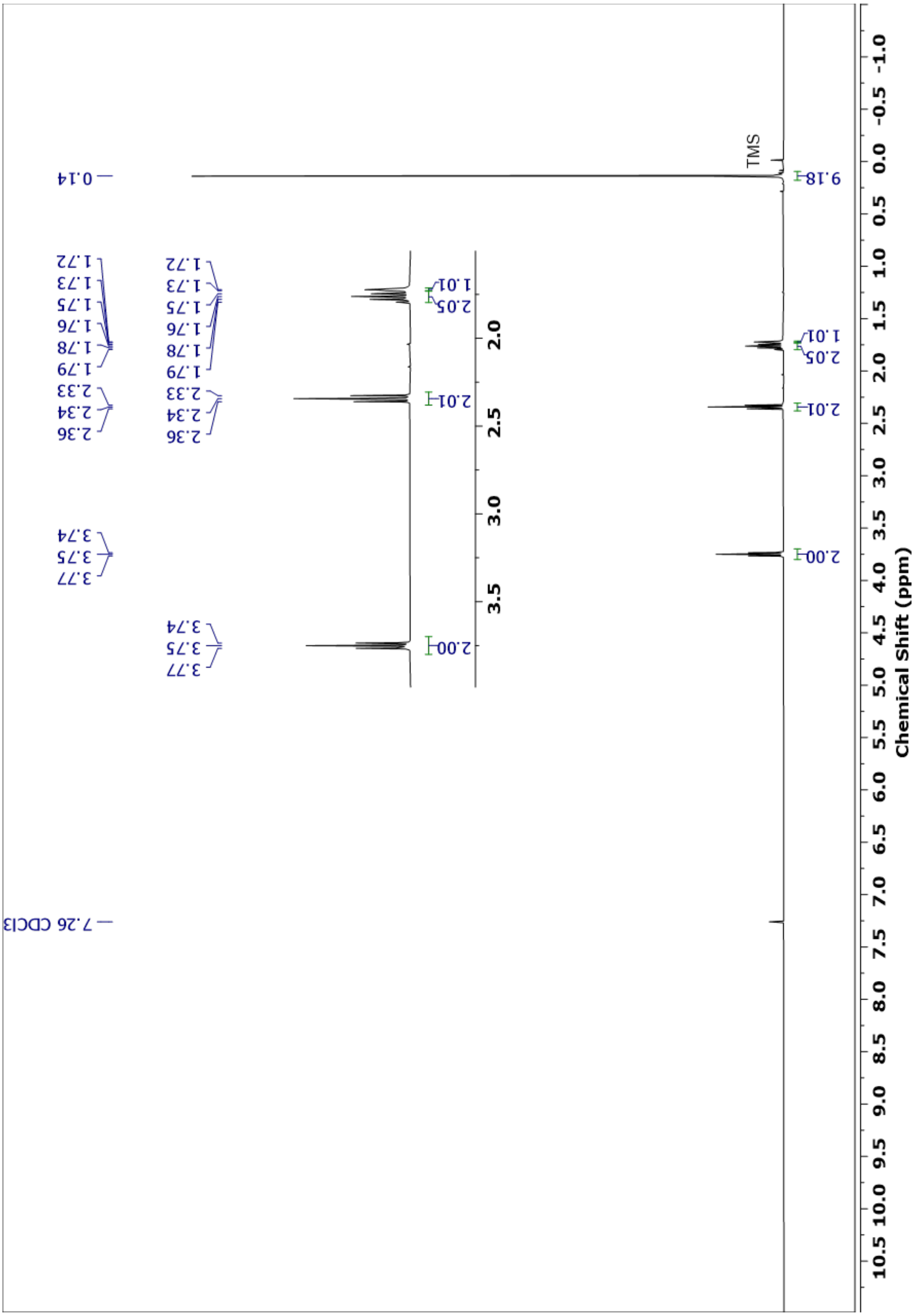

[1]H NMR of (5-Bromopent-1-yn-1-yl)trimethylsilane (2)

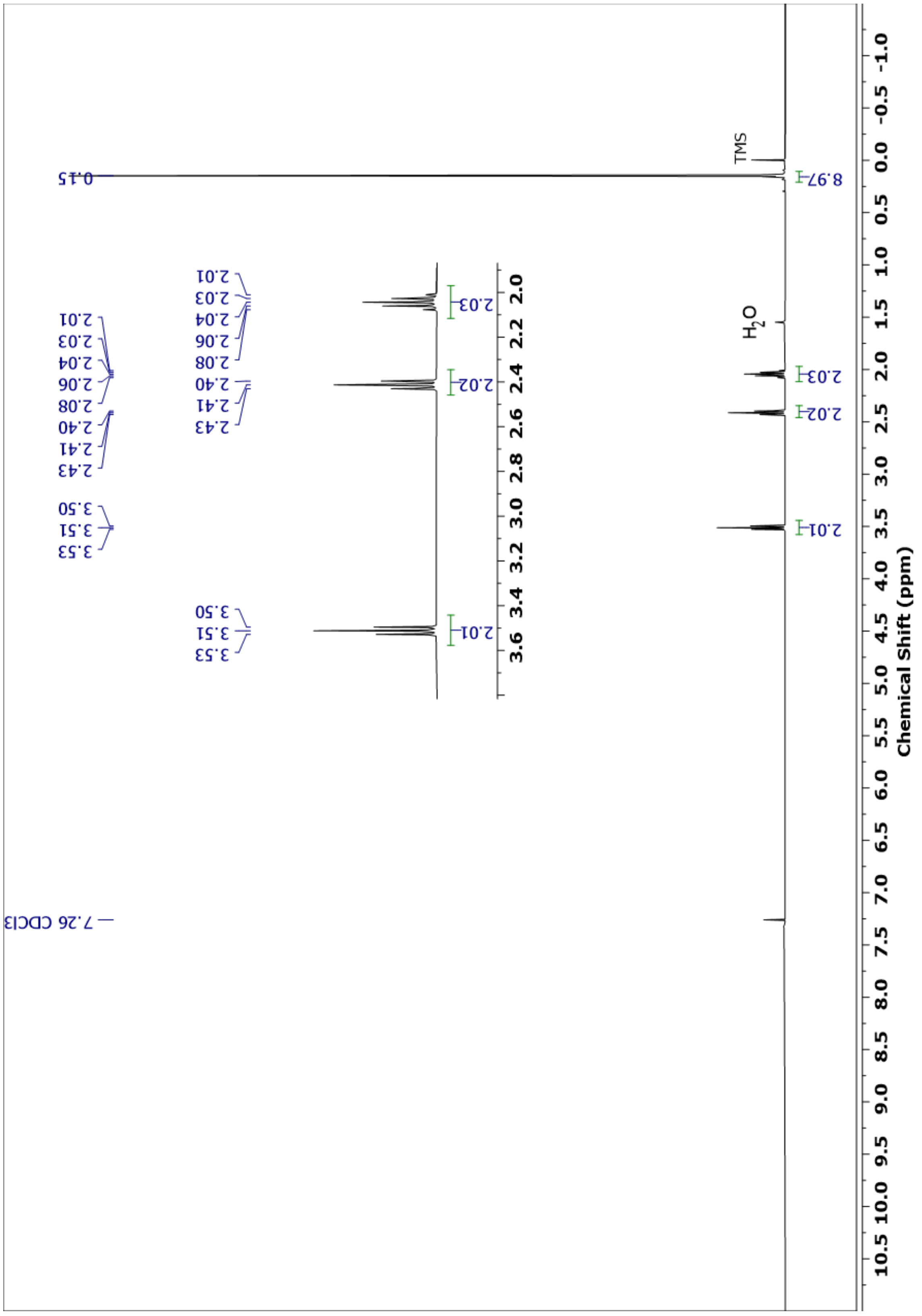

**$^1$H NMR of Methyl 11-oxoundecanoate (4)**

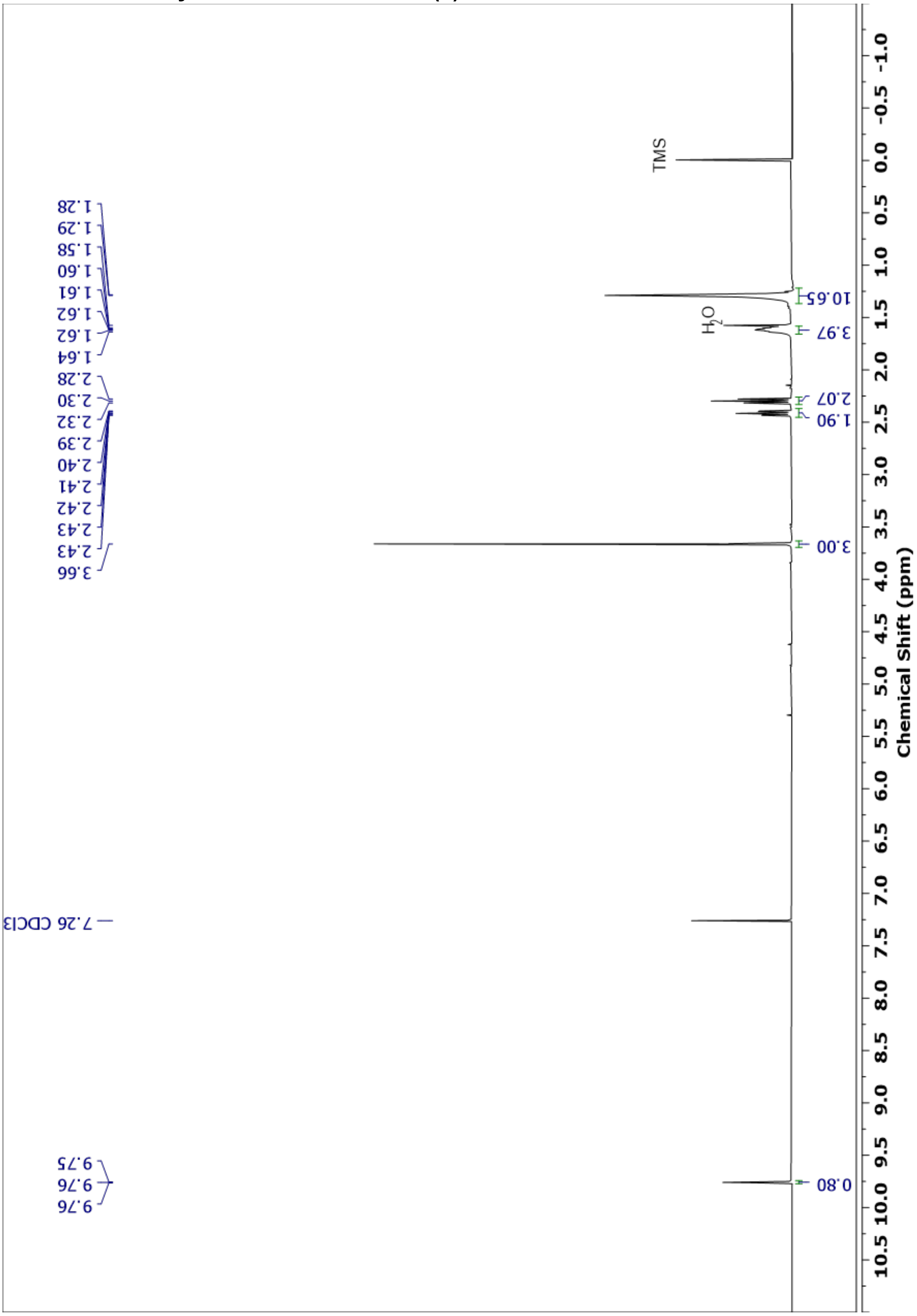

$^{1}$H NMR of Methyl 11-hydroxy-16-(trimethylsilyl)hexadec-15-ynoate (5)

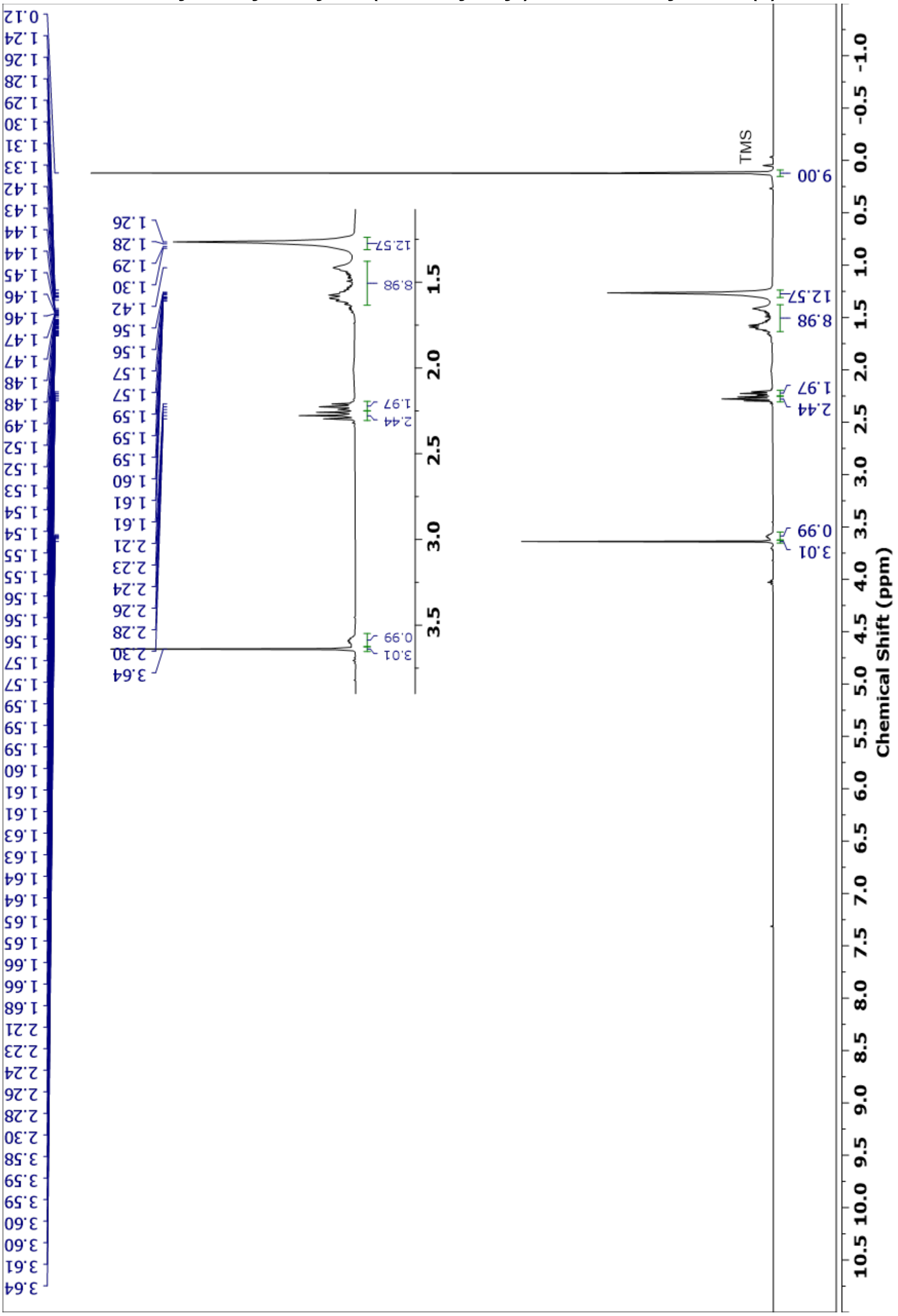

### [1]H NMR of methyl 11-oxo-16-(trimethylsilyl)hexadec-15-ynoate (6)

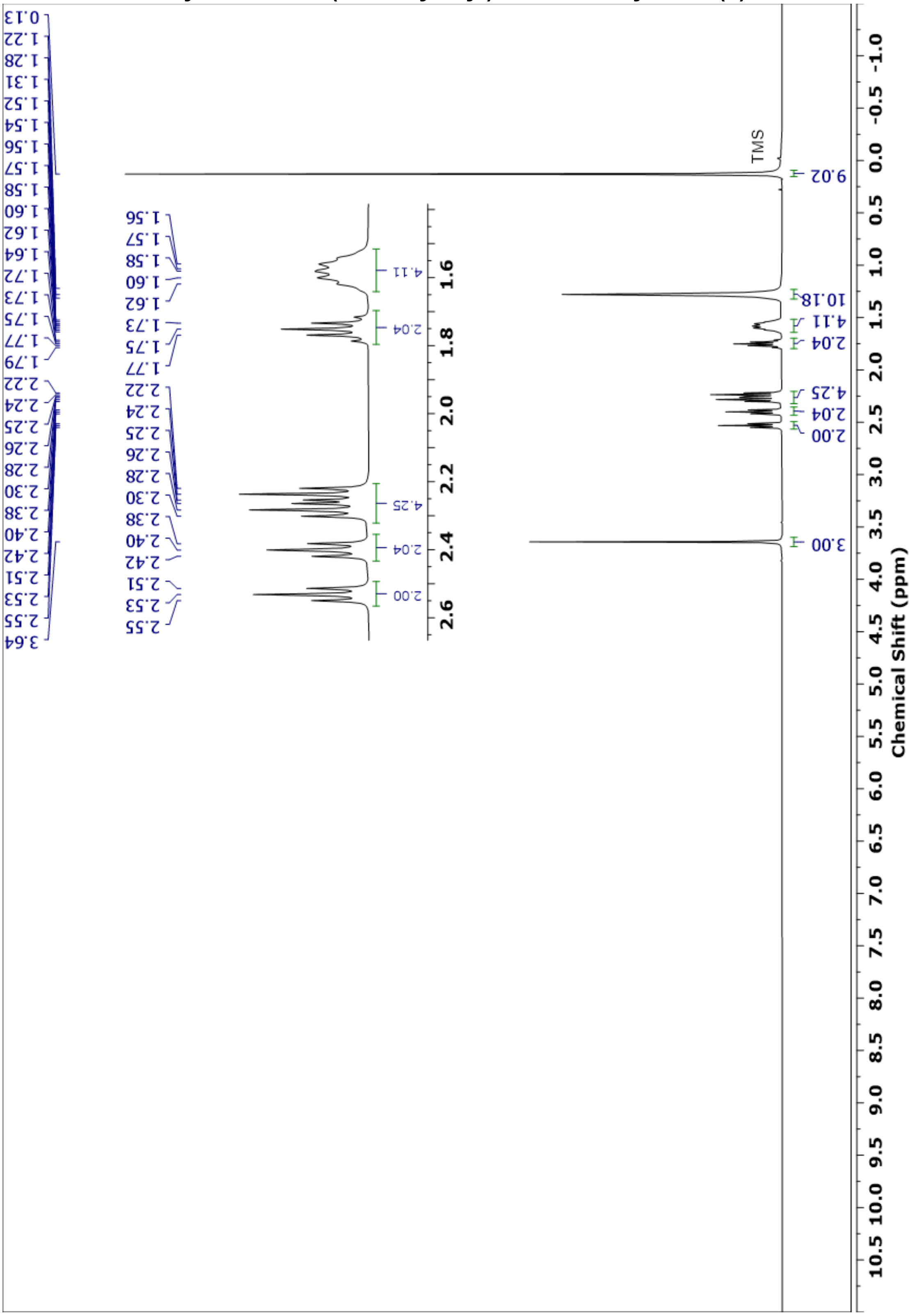

#### $^1$H NMR of 11-Oxohexadec-15-ynoic acid (7)

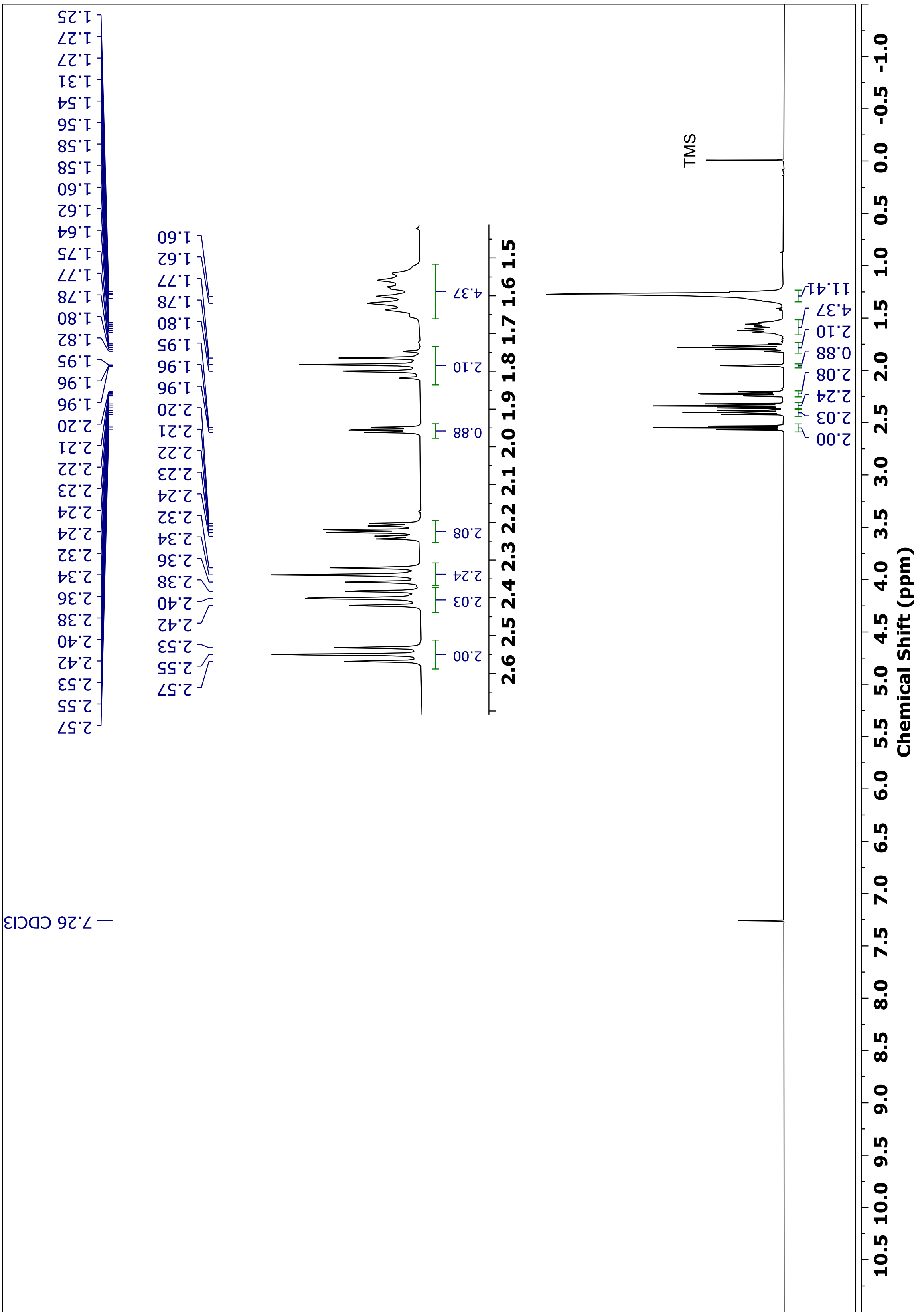

$^{13}$C NMR of 11-Oxohexadec-15-ynoic acid (7)

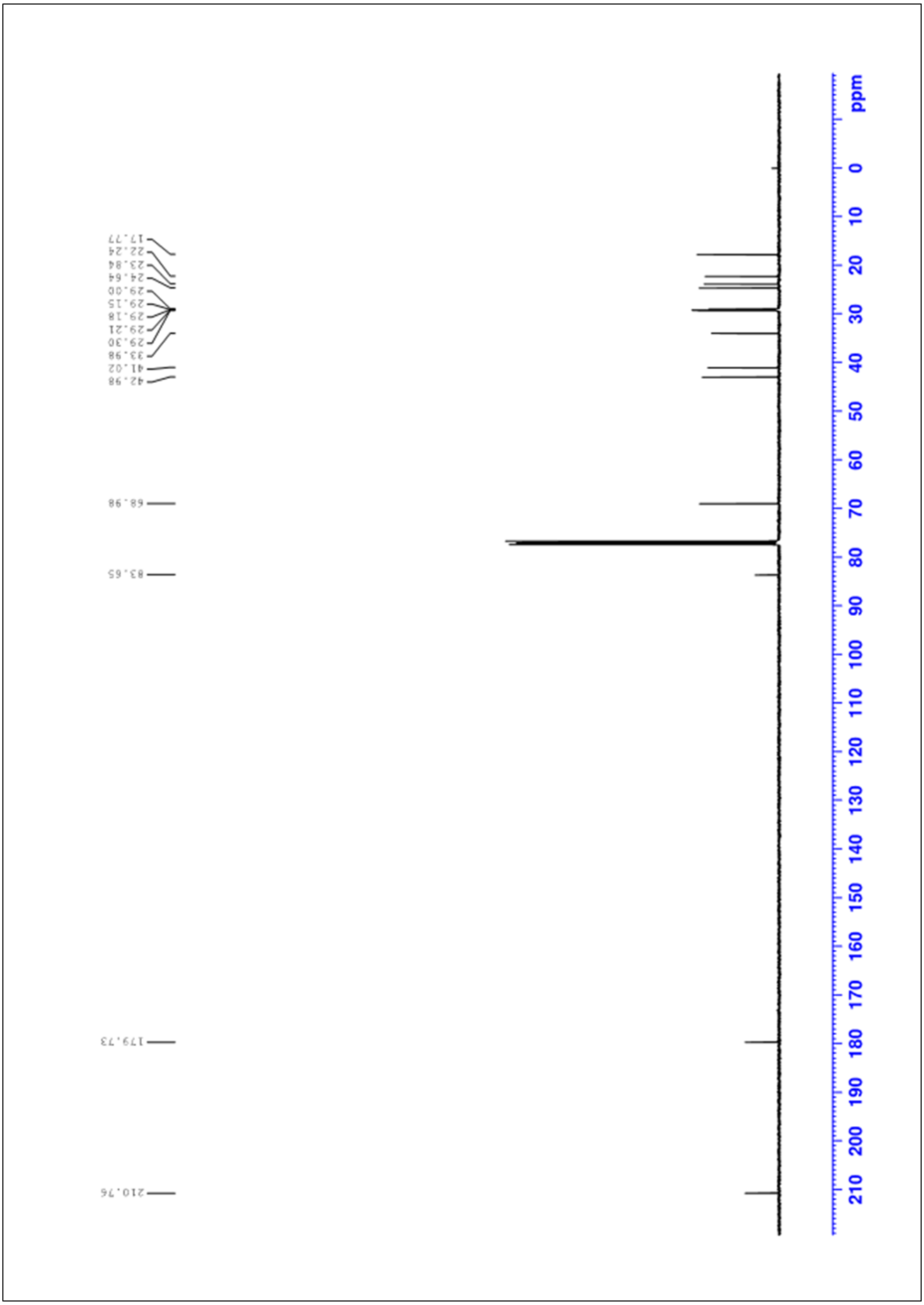

**[1]H NMR of 10-(3-(pent-4-yn-1-yl)-3*H*-diazirin-3-yl)decanoic acid (8) (PA:DA)**

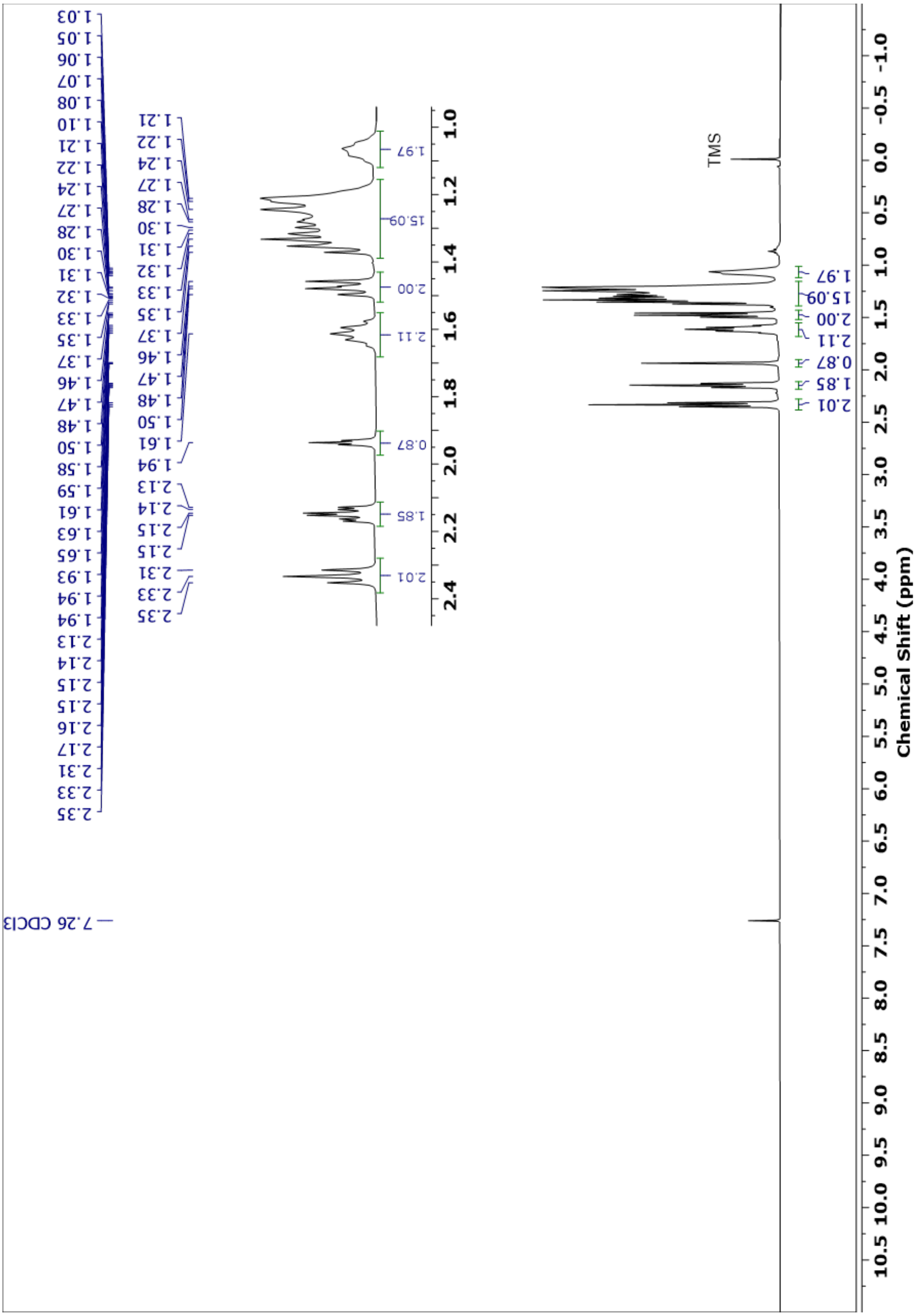

**$^{13}$C NMR of 10-(3-(pent-4-yn-1-yl)-3*H*-diazirin-3-yl)decanoic acid (8) (PA:DA)**

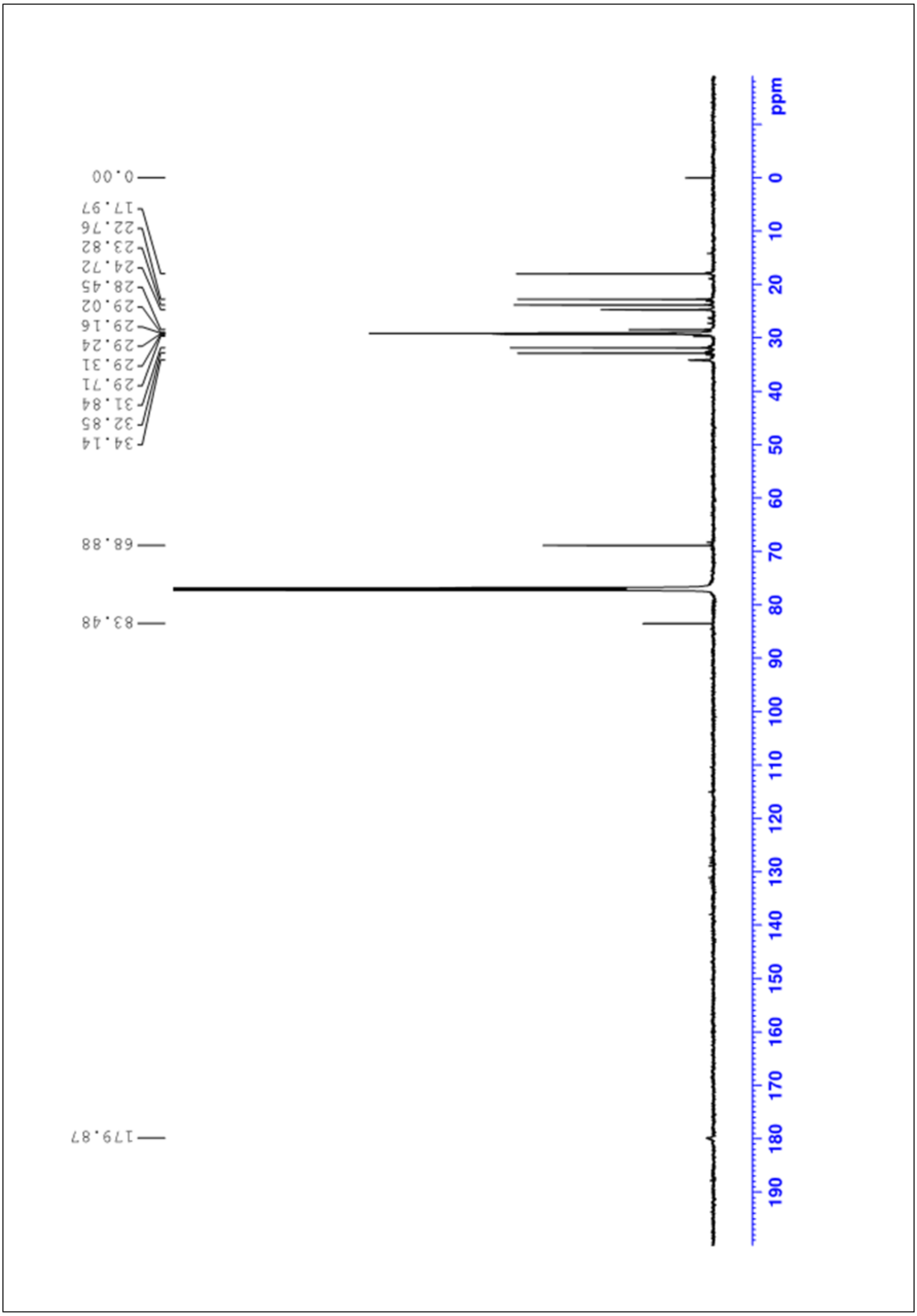

**$^{1}$H NMR of (2,2-Dimethyl-1,3-dioxolan-4-yl)methyl 10-(3-(pent-4-yn-1-yl)-3*H*-diazirin-3-yl)decanoate (9)**

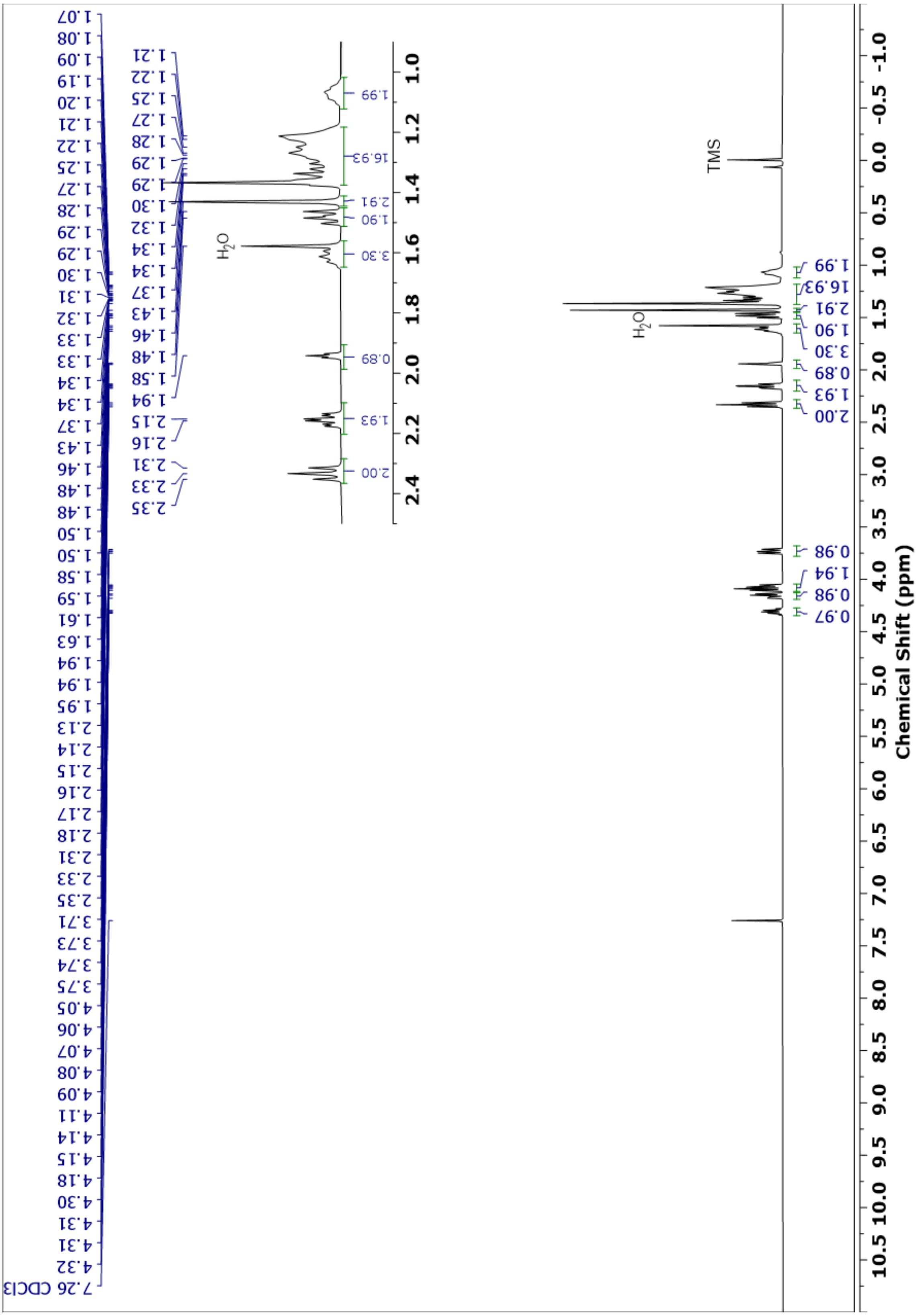

**$^{13}$C NMR of (2,2-Dimethyl-1,3-dioxolan-4-yl)methyl 10-(3-(pent-4-yn-1-yl)-3*H*-diazirin-3-yl)decanoate (9)**

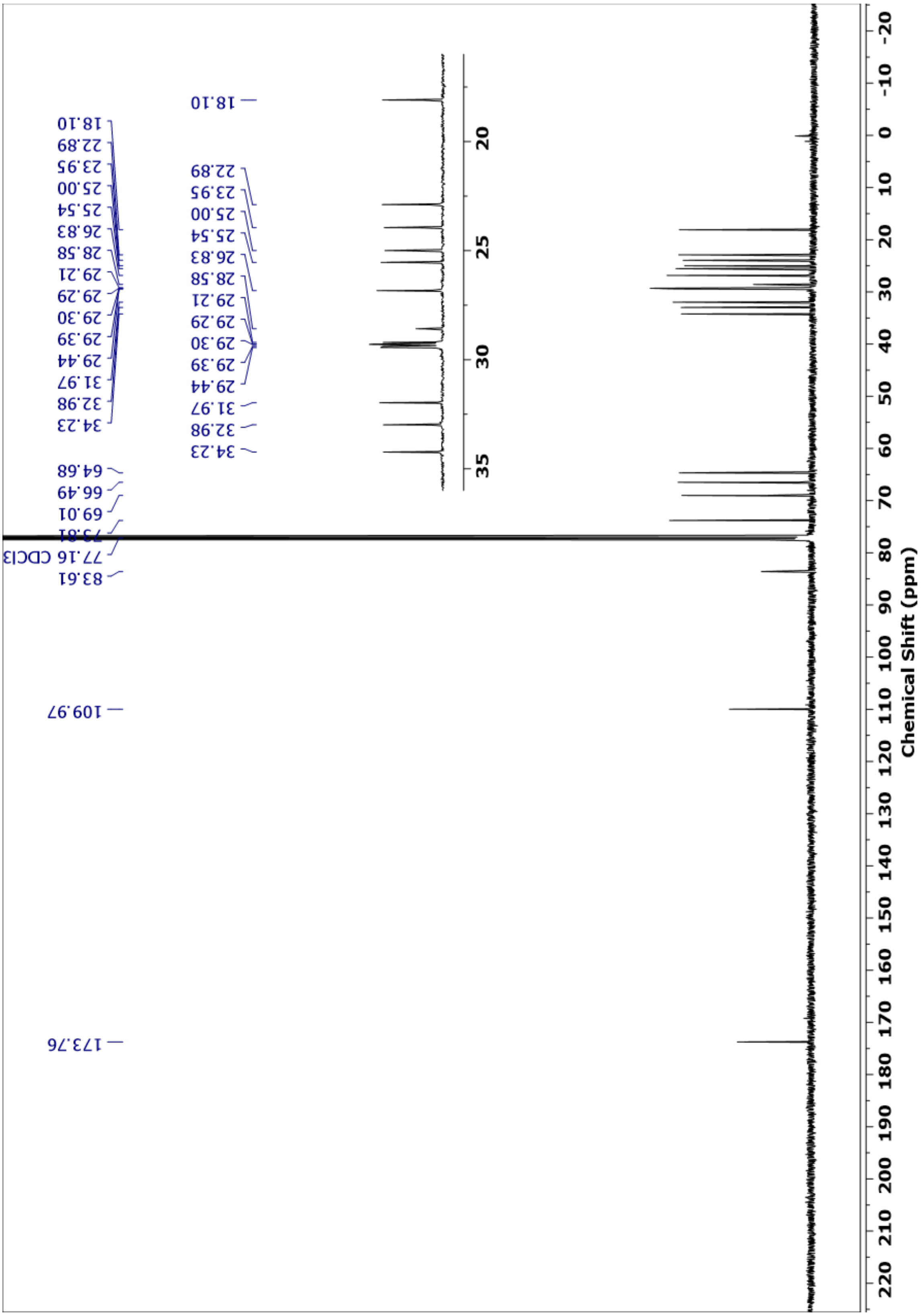

**[1]H NMR of 2,3-dihydroxypropyl 10-(3-(pent-4-yn-1-yl)-3*H*-diazirin-3-yl)decanoate (10) (PG-DA)**

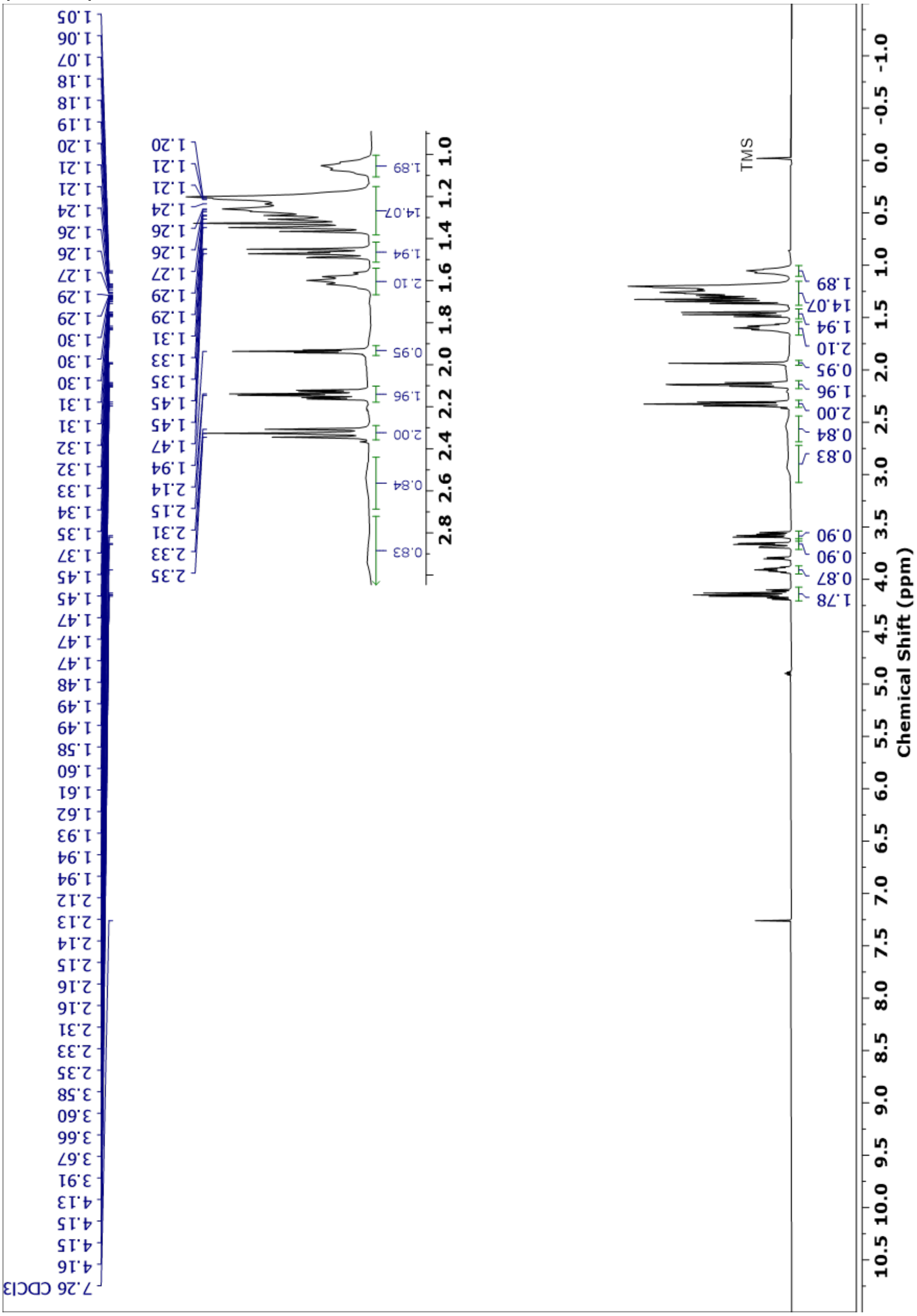

**$^{13}$C NMR of 2,3-dihydroxypropyl 10-(3-(pent-4-yn-1-yl)-3*H*-diazirin-3-yl)decanoate (10) (PG-DA)**

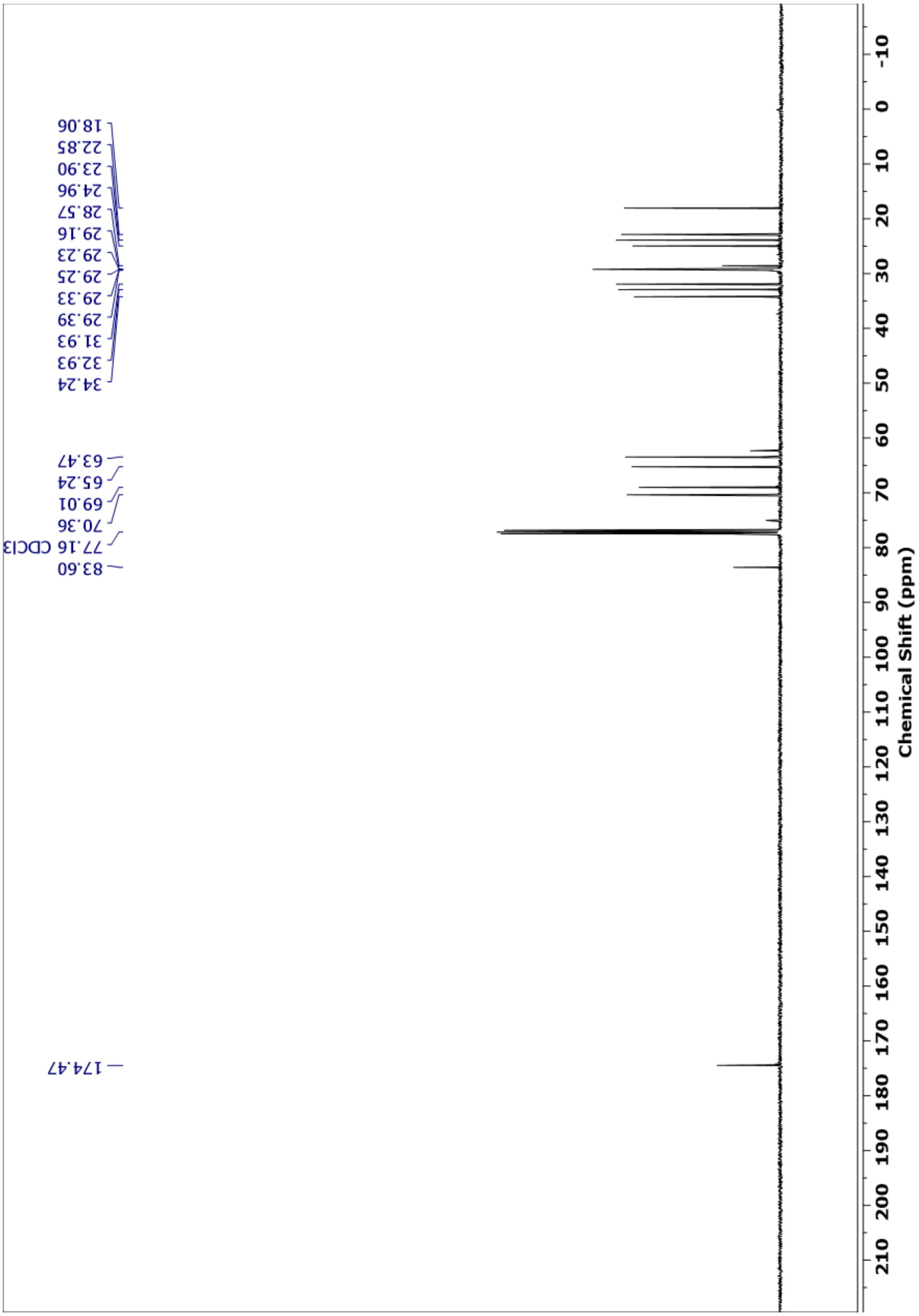
